# Acute exposure to cell-free mitochondrial DNA induces pregnancy-specific aortic endothelial dysfunction and organ-selective inflammation in rats

**DOI:** 10.64898/2026.04.15.718761

**Authors:** Nataliia Hula, Reneé de Nazaré Oliveira da Silva, Desirae Escalera, Leslie Lopez, Gabrielle Kelly, Isabelle K. Gorham, Megan Rowe, Taiming Liu, Arlin B Blood, Eugenia Mata-Greenwood, Xiang-Qun Hu, Lubo Zhang, Nicole R. Phillips, Styliani Goulopoulou

## Abstract

Pregnancy complications such as preeclampsia are associated with circulating cell-free mitochondrial DNA (mtDNA), a damage-associated molecular pattern capable of activating Toll-like receptor 9 (TLR9). We hypothesized that acute mtDNA exposure induces maternal inflammation and endothelial dysfunction during pregnancy via TLR9 activation. Non-pregnant and pregnant rats (gestational days 14-15) were treated intravenously with saline or purified mtDNA and euthanized 4 h after treatment. mtDNA increased cytokine mRNA expression in lung and liver of non-pregnant and pregnant rats, with magnitude varying by pregnancy status and organ. Aortas from pregnant, but not non-pregnant, rats exhibited reduced acetylcholine (ACh)-induced relaxation following mtDNA treatment (Emax, saline: 90.1 ± 3.9 % vs. mtDNA: 62.1 ± 20.7 % KClmax, p<0.05), while uterine artery function was preserved, indicating vascular bed-specific effects. Ex vivo incubation of aortic rings with mtDNA ± white blood cells did not replicate in vivo findings, implicating systemic rather than direct vascular mechanisms. Nuclear DNA did not affect ACh-induced relaxation (p>0.05), confirming that the vascular effects were mtDNA-specific. Pharmacological antagonism of TLR9 with ODN2088 partially attenuated mtDNA-induced maternal endothelial dysfunction. Although overt vascular ROS increases were not detected, aortas from pregnant rats had reduced *sod-1* expression (p<0.05) and increased eNOS protein abundance (p<0.05). Acute mtDNA exposure during pregnancy induces maternal organ inflammation and impairs endothelium-dependent vasodilation, with partial TLR9 involvement. In conclusion, aortic transcriptional changes in antioxidant pathways and increased eNOS abundance were also observed, though their functional significance remains to be determined.

**New & Noteworthy:** To our knowledge, this is the first study to demonstrate that acute exposure to circulating mtDNA induces pregnancy-specific maternal endothelial dysfunction and organ-selective inflammatory responses. Our findings reveal pregnancy- and vascular-bed specific responses of the maternal vasculature to mitochondrial danger signals, with partial TLR9 involvement. Aortic transcriptional changes in antioxidant pathways and increased nitric oxide synthase abundance were identified as molecular correlates of this dysfunction.

## Introduction

Pregnancy is a critical period of immune adaptation characterized by tightly regulated co-operative interactions between the maternal, fetal, and placental immune systems. These dynamic immunological and inflammatory shifts support key pregnancy stages such as implantation, placental development, fetal growth, labor and delivery, while preserving the mother’s ability to respond to exogenous pathogens (1, 2). During pregnancy, the maternal immune system undergoes selective tolerance through compartmentalized rather than systemic, immune modulation, which accounts for greater maternal susceptibility to specific pathogens (3). As such, exposure to certain pathogen-associated molecular patterns (PAMPs; e.g., bacterial infections) or damage-associated molecular patterns (DAMPs; sterile inflammation) can elicit heightened maternal immune responses and are associated with adverse maternal and fetal outcomes (1).

Maternal inflammation and endothelial dysfunction are common pathophysiological features of pregnancy complications (4), such as preeclampsia, and inflammatory signals can promote or exacerbate endothelial dysfunction. Specifically, maternal endothelial cells can undergo functional and morphological changes upon their interaction with circulating inflammatory molecules, resulting in endothelial dysfunction. Endothelial activation can be triggered by bacterial endotoxins and pro-inflammatory cytokines, as well as by direct activation of pattern recognition receptors (PRRs) expressed on endothelial cells following recognition of PAMPs or DAMPs (reviewed in (5)). In addition, endothelial cells can be activated indirectly through their interaction with immune cells and immune cell-derived mediators, amplifying PRR-driven inflammatory signaling and promoting sustained vascular inflammation (5).

Cell-free mitochondrial DNA (mtDNA) is released into the circulation through normal cellular turnover, but under conditions of cellular stress or tissue injury, it can act as a DAMP. Extracellular mitochondria and mitochondrial components can originate from multiple cellular sources and interact with the immune system through diverse mechanisms (6). mtDNA is able to promote immune activation via PRRs such as Toll-like receptor 9 (TLR9) (reviewed in (7, 8)), which recognizes unmethylated CpG motifs enriched in mtDNA. In pregnancy, increased circulating cell-free mtDNA concentrations have been reported in association with inflammatory states and adverse pregnancy outcomes, supporting its potential role as both a biomarker and a mediator of sterile inflammation in gestational complications (7, 9–13).

Supporting a causal role for mtDNA as an inflammatory stimulus, Collins *et al.* demonstrated that intra-articular injection of purified mtDNA in mice promoted immune cell infiltration and increased production of pro-inflammatory mediators (14). Given that cellular responses to CpG DNA can be mediated by TLR9, TLR9 is a plausible innate immune sensor linking mtDNA exposure to downstream inflammatory signaling (8, 15). Consistent with this concept, previous studies have shown a negative impact of TLR9 stimulation on pregnancy outcomes (reviewed in (16)), with TLR9 agonists inducing fetal demise, preterm loss, maternal hypertension (17, 18), disrupted circadian blood pressure regulation (19), and augmented contractile responses in maternal resistance arteries (20).

However, whether purified cell-free mtDNA itself, rather than synthetic TLR9 agonists, can causally induce maternal vascular dysfunction and inflammation during pregnancy remains unknown. Therefore, in the current study, we aimed to evaluate the impact of cell-free mtDNA on maternal inflammation and vascular function during pregnancy. We hypothesized that acute exposure to circulating mtDNA induces maternal inflammation and endothelial dysfunction during pregnancy via TLR9 activation. To test this hypothesis, we acutely challenged pregnant and non-pregnant rats with purified mtDNA with and without pharmacological inhibition of TLR9 and assessed vascular function and inflammatory outcomes. We focused on the thoracic aorta as a model conduit vessel, with uterine arteries included as a pregnancy-adapted reference vascular bed to assess vascular bed specificity of responses. We used a single acute mtDNA exposure to capture early innate immune-mediated responses, consistent with the 3-6 hour window commonly used to assess DAMP-induced inflammatory responses in rodent models (19, 21).

## Materials and Methods

### Chemicals and Reagents

Details of chemicals and reagents used in this study are provided in Supplementary Materials **(Table S1)**.

### Animals

Animal experiments were approved by the Institutional Animal Care and Use Committee (IACUC 22-003) of Loma Linda University, and all procedures were performed in accordance with the Guide for the Care and Use of Laboratory Animals of the National Institutes of Health. The data presented herein are part of a larger research project evaluating maternal vascular and fetoplacental responses to circulating mtDNA. Vascular outcomes reported here were derived from cohorts that overlap with animals from which placental tissue was also collected at the same terminal endpoint for assessments reported separately.

Male, virgin female, and timed-pregnant Sprague-Dawley rats were purchased from Envigo (Indianapolis, IN). On arrival, males were ∼400 g and 13-15-weeks old and virgin females were 190-230 g and 9-11-week-old. Purchased timed-pregnant rats arrived on gestational day (GD)5-8 (term=22-23 days) and were ∼220 g and 11-12-week-old. Rats were pair-housed under 12:12-h light-dark cycle (lights on, 07:00 h; lights off, 19:00 h) in a temperature- and humidity-controlled environment and had free access to tap water and standard laboratory rodent chow. Rats were allowed to acclimate to housing conditions at Loma Linda University animal facilities for one week before handling and experimentation.

In addition to purchasing timed-pregnant rats (Envigo), we bred rats in-house as previously described (19, 22). Briefly, female rats were pair-mated overnight, and the presence of spermatozoa in vaginal smears the following morning was used to designate GD1. Experiments were performed when rats were 12-24-weeks old.

Pregnant rats were studied on GD14-15, with age-matched non-pregnant females used as controls. This gestational age was selected for three reasons. First, earlier exposure to an immune stimulus such as mtDNA could affect placental development and pregnancy viability, which were not primary foci of this study and have been shown to be adversely affected by TLR9 activation at earlier gestational ages (18, 23). Second, GD14-15 represents a gestational period after which fetal growth acceleration is accompanied by substantial maternal cardiovascular adaptations, making it physiologically relevant window for studying maternal vascular responses (24, 25).

Third, this gestational age is consistent with our previous work demonstrating that acute exposure to an innate immune stimulus at comparable gestational ages elicits placental inflammation and disruption of maternal blood pressure circadian rhythms, enabling cross-study comparison of maternal responses (19).

In total, 135 female and 6 male rats were used for this project. Sample sizes for each experiment were based on prior studies and our published protocols (20, 22, 26–29). Male rats were only used for breeding purposes.

### Experimental design and timeline

Three studies were conducted using separate cohorts. <u>Study 1</u> evaluated the in vivo effects of mtDNA on organ inflammation and vascular function in pregnant and non-pregnant rats. <u>Study 2</u> examined whether mtDNA directly affects the vascular function by testing the ex vivo effects of mtDNA on isolated arteries. <u>Study 3</u> investigated the contribution of TLR9 signaling to mtDNA-induced vascular dysfunction through in vivo pharmacological antagonism of TLR9.

Purified DNA mtDNA and nuclear DNA (nDNA) were isolated from liver of GD14 pregnant rats and age-matched non-pregnant rats. The liver was selected as the mtDNA source because hepatocytes are among the most mitochondria-rich cell types in mammals, providing high-yield mtDNA preparations, and because liver-derived mitochondrial DAMPs have been used in established rodent models of systemic inflammatory responses analogous to the current experimental design (21, 30). Donor pregnancy status was matched to recipient status to maximize the physiological relevance of the mtDNA preparation to the inflammatory context being modeled.

Tissue inflammation was measured in lung and liver using reverse transcription quantitative polymerase chain reaction (RT-qPCR), as these organs represent key sites of innate immune activation in response to circulating DAMPs (21); the liver via sinusoidal endothelial cell- and resident Kupffer cell-mediated TLR9 signaling (31, 32) and the lung via its direct exposure to circulating immune stimuli through the pulmonary circulation. Inflammatory gene expression was also assessed in aortic tissue. Vascular function was assessed in isolated arteries using pin and wire myography. Rats were euthanized and tissues were collected 4 h after treatment to capture early, innate immune system-mediated responses while minimizing confounding effects of chronic systemic changes and compensatory mechanisms. Experiments were conducted only in female rats because this study assessed maternal physiology during pregnancy.

### Animal treatments

In Study 1 and Study 3, female rats received a tail-vein injection of sterile 0.9% saline (vehicle) or mtDNA (300 µg/kg body weight). The mtDNA dose was selected based on prior work showing acute systemic inflammatory response in rats at comparable doses (30). Injection volume was adjusted for body weight and mtDNA preparation concentration to achieve the target dose. In Study 1, a subset of animals received nuclear DNA (nDNA, 300 µg/kg body weight) to assess mtDNA specificity. All injections were performed under isoflurane anesthesia (5% for induction, 3% for maintenance, 100% oxygen).

In Study 3, we evaluated the contribution of TLR9 to maternal vascular responses to mtDNA. For this study, female rats received an intravenous injection of either sterile 0.9% saline (Saline group), mtDNA (300 µg/kg body weight; mtDNA group), ODN2088 (TLR9 antagonist, 60 µg/kg body weight) (33) plus 0.9% saline (ODN2088 group), or ODN2088 (60 µg/kg body weight) plus mtDNA (300 µg/kg body weight) (ODN2088+mtDNA group). ODN2088, an inhibitory nucleotide serving as a TLR9 antagonist, was administered 5 min before saline or mtDNA injection. ODN2088 stock solutions (500 mM) were prepared by resuspending lyophilized ODN2088 with endotoxin-free water. The stock solution was then diluted in sterile saline for in vivo treatments.

Before and during treatments, rats were maintained warm using a heating pad. Intravenous injections were administered via the tail vein and delivered within 1 min using a 1 mL syringe fitted with a 25G x 5/8’’ needle. All solutions were prepared and administered under sterile conditions. After treatment, rats were returned to clean cages and were monitored during their recovery. Injections were administered between 8:00 – 9:00 am.

### DNA extraction for in vivo and in vitro treatments

Purified mtDNA (in vivo injections and ex vivo artery incubations) and nDNA (in vivo injections) were isolated from donor rat liver as described below. mtDNA and nDNA were prepared from pregnant and/or age-matched non-pregnant donors as indicated for each experiment.

#### Mitochondrial enrichment

Mitochondria were isolated from liver tissue using the Mitochondria Isolation Kit (Thermo Fisher Scientific) according to the manufacturer’s instructions. The resulting mitochondrial pellet was used for mtDNA extraction.

#### Nuclear enrichment

Nuclei were isolated from liver tissue as previously described (34). Briefly, liver was gently homogenized and passed through a 40-mm cell strainer (Falcon®, Corning, NY, NY, USA, Cat. No. 352340) into ice-cold buffer A (250 mM sucrose, 5 mM MgCl_2_, and 10 mM Tris–HCl, pH 7.4). The homogenate was centrifuged at 600 × g for 10 min at 4 ℃, and the pellet was then resuspended in ice-cold buffer B (2.0 M sucrose, 1 mM MgCl_2_, and 10 mM Tris–HCl, pH 7.4) and centrifuged at 16,000 × g for 30 min at 4 ℃. The resulting nuclear pellet was used for DNA extraction.

#### DNA isolation, quantification, and mtDNA quality control testing

DNA was isolated from mitochondrial and nuclear pellets using the QIAamp DNA Mini Kit (QIAGEN LLC) according to the manufacturer’s instructions. DNA concentration was quantified by spectrophotometry (NanoDrop™ OneC; Thermo Fisher Scientific, Waltham, MA, USA), and DNA was stored at 4 ℃ and used within 1 month after extraction.

Purity and integrity of mtDNA preparations were evaluated by spectrophotometry, qPCR-based assessment of mtDNA enrichment relative to nDNA, protein assay, and TapeStation-based sizing. A260/280 ratios were consistently between 1.8-2.1 qPCR indicated a low level of nDNA carryover in mtDNA preparations; therefore, nDNA injections were included as a specificity control. Protein contamination was assessed using a bicinchoninic acid (BCA) assay and was not detectable in mtDNA preparations. Endotoxin was measured using a chromogenic Limulus amebocyte lysate (LAL) assay (Thermo Fisher Scientific) and was below the assay’s limit of detection in a randomly selected subset of preparations. TapeStation analysis demonstrated a predominant high-molecular weight peak (∼20 kb), consistent with near full-length mtDNA (∼16.3 kb), with limited fragmentation evident as modest peak broadening that varied between samples. Representative quality control data are shown in supplementary materials (**Figure S1**).

### Euthanasia, tissue harvest, and processing

Rats were anesthetized with isoflurane (5% for induction, 3% for maintenance, 100% oxygen) and euthanized by isoflurane overdose followed by bilateral thoracotomy and removal of their hearts. Whole blood was collected from the inferior vena cava under deep anesthesia into EDTA-coated tubes (BD, Franklin Lakes, NJ; Cat No. 367856) and heparin-containing tubes for plasma isolation. After euthanasia, maternal aorta, uterus, lung, and liver were excised and placed in ice-cold Krebs solution of the following composition (in mM): 130 NaCl, 4.7 KCl, 1.18 KH_2_PO_4_, 1.8 MgSO_4_, 14.9 NaHCO_3_, 5.6 Dextrose, 1.56 CaCl_2_·H_2_O. Aortas and uterine arteries were cleaned from perivascular connective and adipose tissue before being used for vascular reactivity experiments. In addition, aortic segments were either stored in Tissue-Tek O.C.T Compound (Sakura, Torrance, CA, USA) for immunofluorescence staining or snap-frozen for further molecular assessment of protein and RNA levels with Western blotting and RT-qPCR, respectively.

Plasma was isolated from whole blood collected in EDTA-coated tubes by centrifugation at 2,000 × g for 15 min at 4 ℃ for subsequent quantification of cell-free mtDNA.

### Vascular function studies

To determine whether mtDNA-induced vascular effects reflect systemic mechanisms or direct actions on the vascular wall, vascular function experiments were performed (1) 4 h after in vivo treatments or (2) following 4 h after ex vivo arterial incubation of arteries with mtDNA or saline.

#### Mounting, normalization, and viability protocols

Isolated thoracic aortas from pregnant and non-pregnant rats were the primary vessels of interest and were used across all vascular studies. Main uterine arteries from pregnant rats were studied as a reference vascular bed to determine whether mtDNA-induced effects were vascular bed specific, given the significant structural and functional remodeling of uterine arteries during pregnancy (35). Accordingly, thoracic aortas and main uterine arteries were used for ex vivo assessment of vascular isometric tension as previously described by Wencelsau et al. with some modifications (36). Briefly, 2-mm aortic rings were mounted in a pin myograph and 2-mm uterine artery rings in a wire myograph, and resting tension was determined (Danish Myo Technology A/S, Aarhus, Denmark). Chambers were filled with 5 mL Krebs-Henseleit solution and continuously gassed with 95% O_2_, 5% CO_2_ at 37 ℃. Optimal baseline tension was then determined for each vascular segment using the DMT Normalization Module for LabChart Software (ADInstruments, Colorado Springs, CO) and vessels were equilibrated for 1 h at baseline tension.

Vascular viability was confirmed by two contractions to high-K^+^ (120 mM KCl). Endothelial integrity was assessed in response to a bolus of acetylcholine (ACh, 3 mM) following pre-constriction with a bolus of phenylephrine (PE, 3 mM).

#### Vascular isometric tension measurements after in vivo mtDNA challenge

For experiments with aorta, cumulative concentration-response curves (CCRCs) were generated for PE (0.1 nM - 30 mM), thromboxane A_2_ receptor agonist U46619 (0.001 nM - 10 mM), ACh (0.1 nM - 30 mM), sodium nitroprusside (SNP, 0.1 nM - 30 mM) and KCl (4.7 - 80 mM). Pre-constriction for ACh and SNP CCRCs were performed with a bolus of PE (1 mM). The force generated during pre-constriction period was similar between groups.

In some aortic rings, the endothelium was mechanically removed (“-E”) by gentle luminal rubbing with a pipette tip. Successful denudation was confirmed in vessels with less than 10% relaxation to ACh. To assess the contribution of NO, vessels were incubated with Nω-Nitro-L-arginine methyl ester hydrochloride (L-NAME, 100 mM) for 30 min (20) before PE pre-constriction and throughout the ACh CCRCs. For uterine arteries, we performed CCRCs to ACh, SNP, and KCl.

#### Vascular measurements after ex vivo incubation with mtDNA

Thoracic aortas from untreated pregnant rats (GD14-15) were placed in a culture dish containing DMEM (supplemented with 1% of charcoal-stripped fetal bovine serum (FBS), 100 U/mL penicillin, 100 𝜇g/mL streptomycin, and L-glutamine) and incubated at 37 ℃ in a humidified incubator for 4 h, as previously described (37).

For treatments, aortas were incubated with saline or mtDNA at various concentrations (0.02, 2, or 20 ng/µL) isolated from pregnant donor liver mitochondria as described above. mtDNA treatment concentrations were selected based on previously reported circulating mtDNA levels in inflammatory conditions (38–40).

To assess the contribution of immune cells to aortic responses to mtDNA, aortic rings were incubated with mtDNA in the presence and absence of autologous white blood cells (WBCs, count: 1 × 10^6^ cells/vessel). Isolation of total WBCs is detailed below. Following incubation, aortas were mounted on a pin myograph. Mounting, normalization, and vascular and endothelial integrity protocols were performed as described above. Vasorelaxation and vasoconstriction were assessed via CCRCs to ACh (0.1 nM - 30 mM; pre-constriction was performed with 60 mM KCl solution) and KCl (4.7 - 80 mM).

For ex vivo incubation experiments, 60 mM KCl was used for pre-constriction rather than PE, because prolonged incubation of isolated vascular segments reduces α_1_-adrenergic receptor-mediated constriction (41). KCl pre-constriction produced comparable contractile responses across all experimental groups.

#### Vascular data analysis and outcomes

CCRCs were analyzed using sigmoidal nonlinear regression (Prism, version 10.0; GraphPad Software Inc., San Diego, CA, USA) and the following parameters were calculated: (i) EC_50_ (expressed as pEC_50_: negative logarithm of EC_50_) and (ii) Emax (maximum response to agonist). The area under the curve (AUC) was calculated from the CCRCs and defined as the cumulative contractile response. Contractile responses are expressed as a percentage of the maximum response to KCl (120 mM).

#### Myography drug dilutions

Stock solutions of PE, ACh, SNP, and L-NAME were prepared in Milli-Q distilled water, whereas U46619 were dissolved in dimethyl sulfoxide (DMSO). Dilutions were prepared fresh on the day of the experiments.

### Biochemical analyses

#### Plasma cell-free mitochondrial DNA quantification

##### DNA extraction

DNA was extracted from approximately 200 µL of plasma using the DNeasy Blood and Tissue kit (QIAGEN) according to the manufacturer’s instructions, starting at the lysis step. For samples with a starting volume < 200 µL, the entire sample was used, and the pre-binding reagents (proteinase K, AL buffer, and 100% ethanol) were adjusted proportionally before loading onto the column. After DNA binding, standard volumes of the remaining reagents were used. DNA was eluted in 200 µL of DNase-free water and concentrated to 20 µL using the Vacufuge vacuum concentrator (Eppendorf).

##### Mitochondrial DNA quantification via qPCR

Quantification of both nDNA and mtDNA was performed through quantitative PCR (qPCR) using the 7500 Real-Time PCR System (Applied Biosystems™, Waltham, MA, USA). The qPCR was performed following the methodology published in Nicklas *et al.* (42) with some modifications, including the omission of the mitochondrial deletion target (26). mtDNA was quantified via the mitochondrial D-loop and nDNA was quantified via the β-actin gene. The qPCR master mix was prepared using 13 µL of TaqManTM Universal Master Mix II, no UNG (Applied Biosystems™) per sample along with the primers and probes listed in **Table S2.** The prepared master mix was added to the wells of a 96-well plate along with either 2 µL of DNA sample or DNase-free water to serve as a negative control. Absolute quantification of the mtDNA target was performed through running a mtDNA standard curve of known mtDNA copy numbers along with each batch of the unknown samples (Sequence: GGTTCTTACTTCAGGGCCATCAATTGGTTCATCGTCCATACGTTCCCCTTAAATAAG ACATCTCGATGGTAACGGGTCTAATC; Cell-free mtDNA is expressed in copies/mL plasma).

### Circulating nitric oxide metabolite quantification

Plasma was isolated from heparin-containing tubes by centrifugation at 9,600 × g for 30 s for subsequent measurements of circulating nitrate and nitrite concentrations (43). Freshly isolated plasma was snap-frozen in liquid nitrogen and stored at −80 ℃ for further analysis of nitric oxide metabolites.

In this manuscript, the term nitrite refers to nitric oxide-related species detected by the assay, including nitrite, S-nitrosothiols, iron-nitrosyls, and free NO. Plasma nitrite levels were measured using the tri-iodide (I₃⁻) assay in combination with an ozone-based chemiluminescence NO analyzer (280i, Sievers, Boulder, CO, USA), as previously described (43–45). Nitrate was assessed using the NiR + I₃⁻ assay, which detects nitrate in addition to the targets of the I₃⁻ assay. For this method, samples were incubated with NiR (a mixture of nitrate reductase, FAD, and NADPH) at 37 ℃ for 45 min, followed by analysis using the I₃⁻ assay. Nitrite and nitrate concentrations are expressed as µM.

### Extraction of RNA from maternal lung, liver, and aortic tissues and cDNA synthesis

Total RNA was isolated from maternal lung, liver, and aortic tissue using QIAzol® Lysis Reagent (QIAGEN), followed by purification with the miRNeasy Mini Kit (QIAGEN). RNA purity and quantity were assessed by spectrophotometry (NanoDrop™ OneC).

Reverse transcription was performed using Sensiscript RT Kit reagents (QIAGEN) with RiboGuard RNase inhibitor (LGC, Biosearch Technologies) and oligo-dT primers (QIAGEN), as we previously published (19, 46). Reactions were incubated at 37 ℃ for 1 h, and cDNA was stored at -20 ℃ until qRT-PCR experiments.

### Quantitative real-time polymerase chain reaction

Expression of pro-inflammatory cytokines, anti-inflammatory cytokines, immune cell markers, and housekeeping genes in rat aorta, liver and lung tissues was determined using qRT-PCR. Primer sequences are listed in **Table S3.**

qRT-PCR reactions were performed as previously described (19) using iQ SYBR Green Supermix (Bio-Rad) on a CFX96 Real-Time PCR Detection System with CFX Maestro Software v.2.3 (Bio-Rad). Cycling conditions consisted of an initial denaturation at 95 ℃ for 3 min, followed by 40 cycles of 95 ℃ for 10 sec, and 60 ℃ for 1 min. A dissociation melt curve analysis was performed at the end of each run to verify amplification specificity. Target gene expression was normalized to the reference genes *18s* (for aortic tissue) or *gapdh* (for lung and liver tissues), which were selected based on their stable expression across samples. Relative gene expression was calculated using the 2^-ΔΔCT^ method.

### White blood cell isolation from whole blood

White blood cells (WBCs) were isolated and immediately used for ex vivo artery incubations. Briefly, 10 mL aliquots of whole blood were subjected to red blood cell lysis buffer (per 1L: 155 mM NH4Cl, 10 mM KHCO3, 0.2 mL of 0.1 M EDTA diluted in distilled water; pH: 7.2-7.4; sterilized through a 0.2 µm filter) and incubated for 5 min in 37 ℃ water bath. Cells were centrifuged at 400 × g for 5 min, and the pellet was washed and resuspended in 1× RPMI solution containing 10% FBS (Gibco) and 1× antibiotic-antimycotic (Gibco). Cell numbers were determined using trypan blue (Sigma) exclusion, and 1 × 10^6^ cells per aortic segment were used for ex vivo vessel incubation studies.

### Western blot analysis

Western blot analysis was performed in rat aortic tissue, as previously described (26, 47). Proteins were extracted using T-PER Tissue Protein Extraction Reagent (Thermo Fisher Scientific) supplemented with protease inhibitor cocktail tablets (Sigma Aldrich), and concentrations were determined using a BCA assay. Samples were denatured with 𝛽-mercaptoethanol (Sigma Aldrich) and equal amounts of protein were separated by SDS-PAGE (5 μg for SOD-1; 20 μg for SOD-2, Catalase, eNOS). Proteins were transferred to nitrocellulose or polyvinylidene difluoride (PVDF) membranes using a Trans-Blot Turbo Transfer System (Bio-Rad) or by wet transfer, with membranes and transfer method selected according to the protein of interest (Table S5). Membranes were blocked and incubated with primary and secondary antibodies as detailed in Tables S4-S5.

Protein signals were detected using either Odyssey CLx imaging system (LI-COR Biosciences, Lincoln, NE, USA) or an AZURE 300 system (Biosystems), with analysis performed either with analysis in Image Studio (v5.2) or ImageJ software (v13.0.6, NIH, USA). Signals were normalized to total protein using Ponceau staining or Revert™ 700 Total Protein Stain (LI-COR Biosciences). Data are expressed relative to the saline control group.

### Oxidant- and NO-related fluorescence imaging in rat aorta

Oxidant-sensitive and NO-reactive fluorescence was assessed in thoracic aorta from saline- and mtDNA-treated rats using dihydroethidium (DHE) and (4-Amino-5-Methylamino-2’,7’-Difluorofluorescein Diacetate (DAF-FM) staining, respectively, as previously described (33, 48). Aortic segments were embedded in Tissue-Tek® O.C.T. Compound (Sakura) and snap-frozen in liquid nitrogen. Cross-sections (10 µm) were equilibrated in PBS (3 × 5 min; PBS replaced every 5 min) in a light-protected humidified chamber. For oxidant-sensitive staining, sections were incubated with DHE (10 µmol/L, Sigma-Aldrich) for 30 min. For NO-reactive staining, sections were incubated with DAF-FM Diacetate (20 µmol/L; Invitrogen™, Thermo Fisher Scientific) for 1 h. Negative control sections were processed in parallel using PBS without probe. Slides were imaged on an ECLIPSE Ts2 inverted microscope (Nikon Instruments Inc., NY, USA) using a ×4 objective with TRITC (tetramethylrhodamine isothiocyanate) filter sets for DHE and FITC (fluorescein isothiocyanate) filter sets for DAF-FM. Fluorescence was quantified using ImageJ software (v1.54J). DHE fluorescence was used as an index of oxidant-dependent DHE oxidation within the aortic wall, whereas DAF-FM fluorescence was used as an index of NO-derived nitrosating activity/NO-related signaling following intracellular de-esterification.

### Statistical analysis

Data distributions were assessed using the Shapiro-Wilk test. When data were not normally distributed, log-transformation was applied prior to analysis. Outliers were identified using the ROUT (Robust regression and Outlier) method with Q=1%.

Comparisons were performed using unpaired t-tests for normally distributed data with equal variances, Mann-Whitney *U* tests for non-normally distributed data, one-way ANOVA with Tukey’s post hoc test, or two-way ANOVA with Sidak post hoc test, as appropriate. For RT-qPCR, statistical analyses were performed on ΔCt values. Data are presented as mean ± standard deviation (SD), except concentration-response curves which are presented as mean ± standard error of the mean (SEM). Statistical significance was set at α = 0.05. The specific statistical approach used for each dataset is specified in the corresponding figure legends, and exact p values are reported for all analyses. Analyses were conducted in GraphPad Prism (v10.0.1; GraphPad Software, San Diego, CA, USA).

## Results

### Circulating levels of mtDNA

At euthanasia, plasma cell-free mtDNA was quantified 4 h after injection of saline (control) or mtDNA in pregnant and non-pregnant rats. Plasma mtDNA copy number (copies/mL) was comparable across groups (non-pregnant saline: 497.1 ± 429.4, n=7 rats; non-pregnant mtDNA: 672.5 ± 800.3, n=9; pregnant saline: 315.6 ± 93.8, n=8; pregnant mtDNA: 469.1 ± 350.3, n=9). Statistical analyses were performed on log-transformed data using two-way ANOVA and showed no pregnancy-by-treatment interaction (F(1, 29)=0.0246, p=0.876) and no main effects of pregnancy (F(1, 29)=0.2270, p=0.637) or treatment (F(1, 29)=0.00041, p=0.984). These data indicate that circulating cell-free mtDNA levels were not elevated at 4 h after treatment, consistent with clearance/redistribution by this time point.

### Systemic markers of maternal inflammation

To assess systemic maternal inflammatory activation 4 h following mtDNA exposure, we measured pro-inflammatory cytokines *il-1β, tnf-α, il-6,* and *ifn-γ* and immunomodulatory/anti-inflammatory cytokines *il-10* and *il-4* in liver and lung tissues from non-pregnant and pregnant rats. In addition, we measured *tlr9* expression in these tissues to determine whether innate immune sensing pathways were engaged.

In the liver (**Figure 1A-G, left panels)**, mtDNA exposure elicited a robust inflammatory transcriptional response in both non-pregnant and pregnant rats, with increased expression of *il-1β, il-6, tnf-α, ifn-γ,* and *il-4* (two-way ANOVA with Sidak multiple comparisons test; all p ≤ 0.04). There were significant pregnancy-by-treatment interactions for *il-1β* (F(1, 23)=9.919, p=0.0045), *il-6* (F(1, 23)=6.832, p=0.0156), and *il-10* (F(1, 25)=7.8, p=0.01), indicating that the mtDNA effects differ by pregnancy status. Sidak tests demonstrated that the mtDNA-induced increase in *il-1β* was greater in non-pregnant rats compared with pregnant rats (predicted least squares means diff ± SEM, non-pregnant: 0.598±0.066; pregnant: 0.3001 ± 0.068, p=0.0004), whereas the increase in *il-6* was greater in pregnant rats (non-pregnant: 2.06 ± 0.61, p=0.0051; and pregnant: 4.34 ± 0.63, p<0.0001). Liver *il-10* increased in non-pregnant but did not change in pregnant rats in response to mtDNA (predicted least squares means difference ± SEM, non-pregnant: 0.14 ± 0.027, p<0.001; pregnant: 0.0250 ± 0.03, p=0.65).

**Figure 1.**
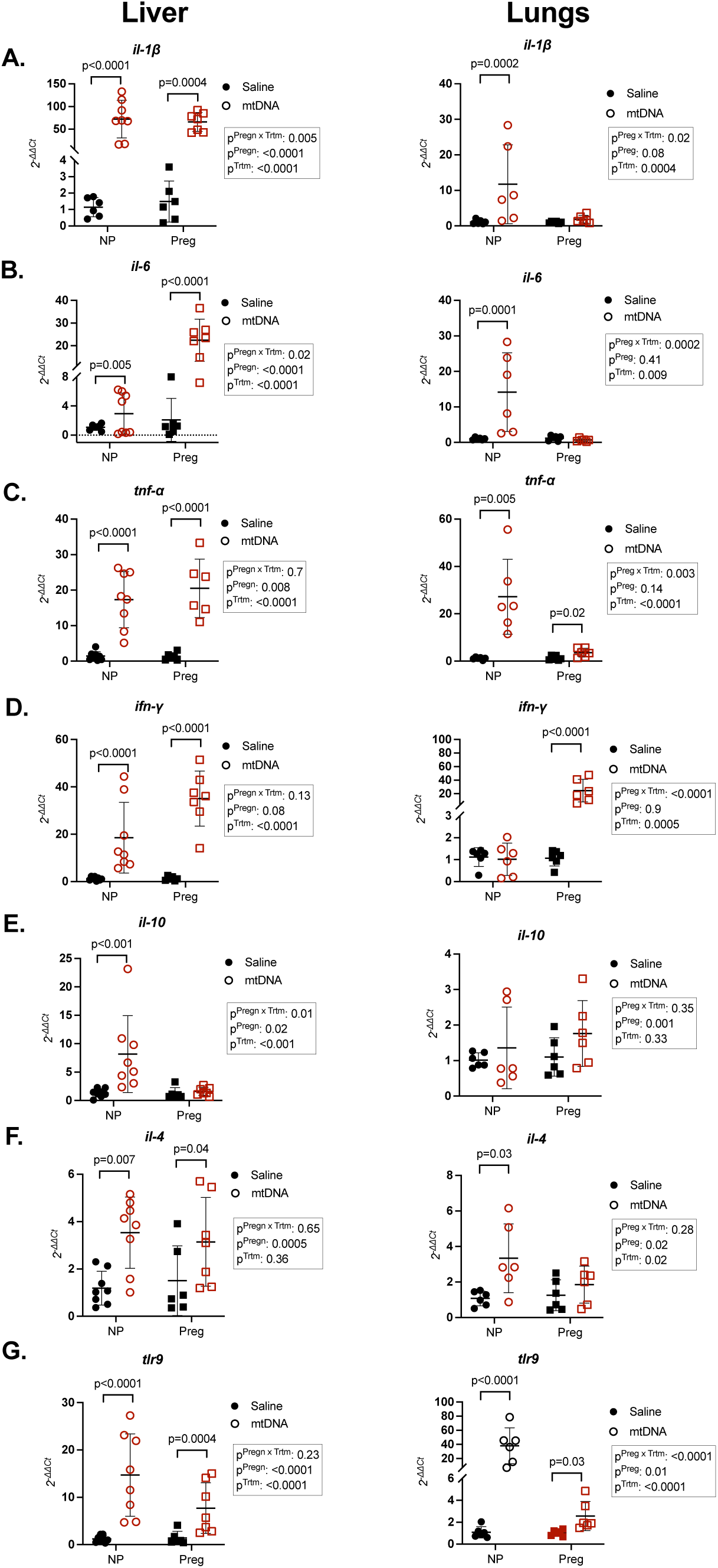
Liver and lung cytokine mRNA expression 4 hours after mitochondrial DNA (mtDNA) treatment. Relative expression (2^-ΔΔCt^) of (A) interleukin (il) 1β *(il-1β)*, (B) tumor necrosis factor α (*tnf-α*), (C) *il-6,* (D) interferon γ (*ifn-γ*), (E) *il-10*, (F) *il-4,* and (G) toll-like receptor 9 (*tlr9*) in liver (left panels) and lung (right panels) from non-pregnant (NP) and pregnant (Preg) rats 4 h after intravenous injection of saline (control) or purified mtDNA. Statistical tests were performed on ΔCt values. Data were analyzed by two-way ANOVA with Sidak post hoc test; individual data points are shown, with bars representing mean ± SD; n=6 rats/group.

In the lung **(Figure 1A-G, right panels)**, mtDNA exposure induced a pregnancy-dependent inflammatory transcriptional response. Two-way ANOVA showed significant (all p≤0.02) pregnancy-by-treatment interactions for *il-1β* (F(1, 20)=6.173, p=0.0219)*, tnf-α* (F(1, 20)=11.16, p=0.0033)*, il-6* (F(1, 20)=21.41, p=0.0002), indicating that the pulmonary cytokine response to mtDNA differ by pregnancy status. Sidak post hoc tests revealed that *il-1β* increased only in non-pregnant rats (p=0.0002), *ifn-γ* increased only in pregnant rats (p<0.0001), whereas *il-6* increased in non-pregnant rats (p=0.0019) and decreased in pregnant rats (p=0.0286). The mtDNA-induced increase in *tnf-α* was greater in the non-pregnant group compared to the pregnant group (mean diff ± SEM, non-pregnant: 4.579 ± 0.543, p<0.0001; pregnant: 1.659 ± 0.543, p=0.0124). Interestingly, *ifn-γ* was the only cytokine that was increased only in lungs of pregnant rats (F(1, 20)=26.19, p<0.0001). There were significant main effects for treatment (F(1, 20)=6.943, p=0.0159) and pregnancy (F(1, 20)=6.071, p=0.0229) for *il-4,* while *il-10* remained unchanged (p>0.05).

mtDNA increased *tlr9* mRNA in both liver **(Figure 1G, left panel)** and lung **(Figure 1G, right panel)** in non-pregnant (p<0.0001) and pregnant (p=0.03) rats, consistent with engagement of innate immune sensing in response to mtDNA exposure. Two-way ANOVA revealed a significant pregnancy-by-treatment interaction for lung *tlr9* (F(1,20)=31.20, p<0.0001), indicating that mtDNA effects on *tlr9* expression in this tissue differ by pregnancy status. Post hoc Sidak tests showed that the mtDNA-induced increase in lung *tlr9* was greater in non-pregnant compared to pregnant rats (predicted least squares means difference ± SEM, non-pregnant: 4.876 ± 0.4626, p<0.0001; pregnant: 1.222 ± 0.462, p=0.031). This pregnancy-dependent pattern in lung *tlr9* expression was consistent with the pregnancy-specific differences observed in pulmonary cytokine response to mtDNA. Together, these data indicate that mtDNA elicited an inflammatory response, as manifested by maternal liver and lung cytokine expression changes, and induced *tlr9* transcription, with the magnitude of these responses varying with pregnancy status and differing across maternal organs.

### Vascular function

Given the systemic inflammatory activation and possible TLR9 engagement we observed, and the association between endothelial function and inflammation documented in previous investigations (7, 9–13), we next tested whether mtDNA exposure alters vascular reactivity, with emphasis on endothelial function.

#### *In vivo* mtDNA challenge – contractile responses

To evaluate changes in aortic contractile function due to mtDNA exposure, we performed CCRCs to PE **(Figure 2A-F)** and U46619 **(Figure 2G-L)** in endothelium denuded (-E) or intact (+E) aortic rings in non-pregnant and pregnant rats at 4 h after injection. Two-way ANOVA demonstrated a main effect of denudation for PE pEC_50_ in both non-pregnant and pregnant rats (all p≤0.05; **Figure 2C, F)**, with no treatment-by-denudation interaction for either constrictor (all p≥0.05). In non-pregnant rats, there was also a main effect of treatment for U46619 (p=0.047; **Figure 2I**). In Sidak-adjusted post hoc comparisons, denudation increased PE pEC_50_ in both saline- and mtDNA-treated non-pregnant groups (all p<0.001; **Figure 2A-C)** and in saline-treated pregnant rats (p=0.05; **Figure 2D, F)**, whereas this comparison was not significant in mtDNA-treated rats (p>0.05; **Figure 2E, F**). Aortic contractile responses to non-receptor-mediated depolarization (KCl CCRC) were comparable between saline-treated and mtDNA-treated rats in both the non-pregnant (p=0.92) and pregnant (p=0.98) groups (two-way ANOVA, n=5-7 rats/group; **Figure S2)**. Detailed outcomes from statistical assessment of the CCRC variables are presented in **Table S6.** Collectively, these data indicate that mtDNA does not alter overall aortic contractile responses but attenuates endothelial modulation of PE-induced constriction in pregnancy.

**Figure 2.**
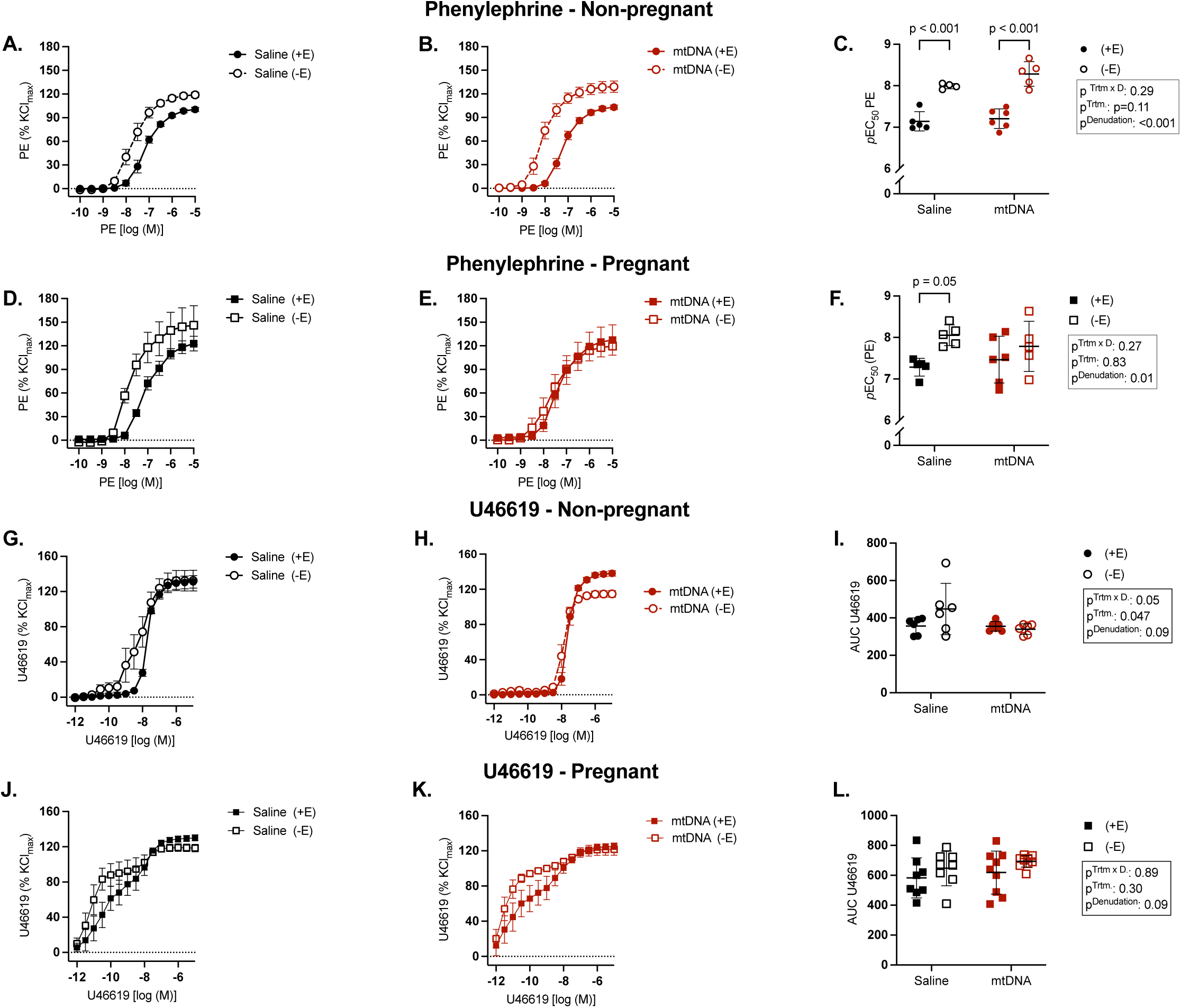
Aortic contractile responses in non-pregnant and pregnant rats after in vivo saline or mitochondrial DNA (mtDNA) challenge. Cumulative concentration-response curves (CCRCs) and corresponding pEC_50_ values for phenylephrine (PE; A-F) and U46619 (G-L) were assessed in thoracic aortic rings from non-pregnant (A-C, G-I) and pregnant rats (D-F, J-L) 4 h after intravenous injection of saline (black symbols) or mtDNA (red symbols). Endothelium-denuded aorta assays are shown in open symbols, saline-treated in black symbols, and mtDNA-treated in red symbols. pEC_50_ is the negative logarithm of the EC_50_, and area under the curve (AUC) reflects total contractile activity across the concentration range. Data were analyzed by two-way ANOVA with Sidak post hoc test. CCRCs are presented as mean ± SEM. pEC_50_ values are shown as individual data points with bars representing mean ± SD; n=6-9 rats/group.

#### *In vivo* mtDNA challenge – endothelium-dependent relaxation responses

To determine whether mtDNA affects endothelium-dependent relaxations, we assessed aortic relaxation responses to ACh (**Figure 3A-C**). Two-way ANOVA showed a main effect of pregnancy (F(1, 21)=15.36, p=0.0008), a main effect of treatment (F(1, 21)=10.57, p=0.004), and a significant pregnancy-by-treatment interaction (F(1, 21)=5.410, p=0.03) for ACh E_max_. Sidak adjusted post hoc analysis showed that mtDNA treatment reduced E_max_ in pregnant (p=0.0017; n=6-8 rats/group) but not in non-pregnant rats (p=0.96; n=5-6 rats/group). There were no differences between non-pregnant and pregnant saline rats in relaxation responses (E_max_, p=0.76); however, in the presence of mtDNA, pregnant rats had lower relaxation responses compared to non-pregnant rats (E_max_, p=0.0005). A similar pattern was observed for ACh pEC_50_ **(Table S6)**. These results indicate that mtDNA impairs endothelium-dependent relaxations in the aorta in a pregnancy-specific manner.

**Figure 3.**
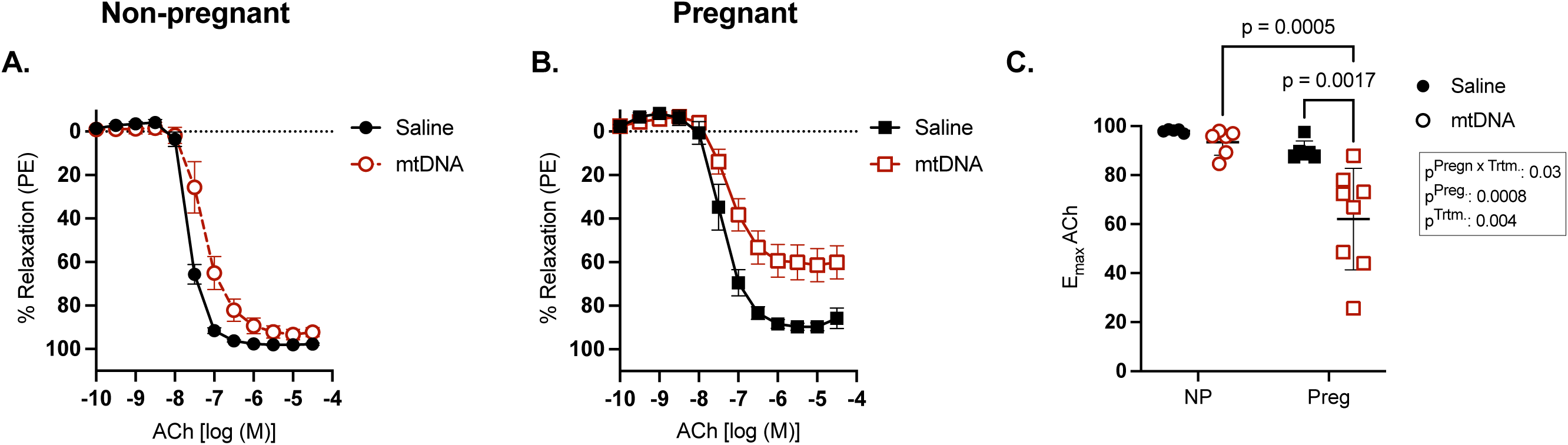
Aortic endothelium-dependent relaxation responses in non-pregnant and pregnant rats after in vivo saline or mitochondrial DNA (mtDNA) challenge. Cumulative concentration-response curves (CCRCs) and corresponding maximum responses (E_max_) for acetylcholine (ACh) were assessed in thoracic aortic rings from in non-pregnant (A, C) and pregnant (B, C) rats 4 h after intravenous injection of saline (black symbols) or mtDNA (red symbols). Data were analyzed by two-way ANOVA with Sidak post hoc test. CCRCs are presented as mean ± SEM. E_max_ values are shown as individual data points with bars representing mean ± SD; n=5-8 rats/group.

#### In vivo nDNA challenge – treatment specificity

Next, we assessed whether the mtDNA-induced vascular effects observed in pregnant rats were specific to mtDNA. In this set of experiments, we measured PE-induced contractile responses and ACh-induced dilatory responses in aortas from pregnant rats 4 h after intravenous injection of nDNA or saline. For these comparisons, the saline control group was derived from the corresponding saline dataset shown in the prior PE and ACh experiments **(Figures 2, 3).** Two-way ANOVA showed no significant main effects or interaction for PE E_max_ (p≥0.17; **Figure 4A, B**). There were significant effects of denudation (F(1,14)=35, p<0.001) and treatment (F(1, 14)=9.1, p=0.009) for PE pEC_50_, with no denudation-by-treatment interaction (F(1, 14)=0.18, p=0.68) **(Figure 4A, C)**. Sidak-adjusted paired comparisons showed that denuded aortas from both saline and nDNA treated rats had greater pEC_50_ compared with endothelium intact vessels (all p£0.004). In addition, ACh responses were similar between saline and nDNA groups, with no differences in E_max_ (p=0.61; Mann-Whitney *U* test) or pEC_50_ (p=0.32; unpaired t-test; n=4-6 rats/group; **Figure 4D-F**). Together, these data show that nDNA does not alter aortic contractile and dilatory functions in pregnant rats, suggesting that the endothelial dysfunction observed in this study was specific to mtDNA.

**Figure 4.**
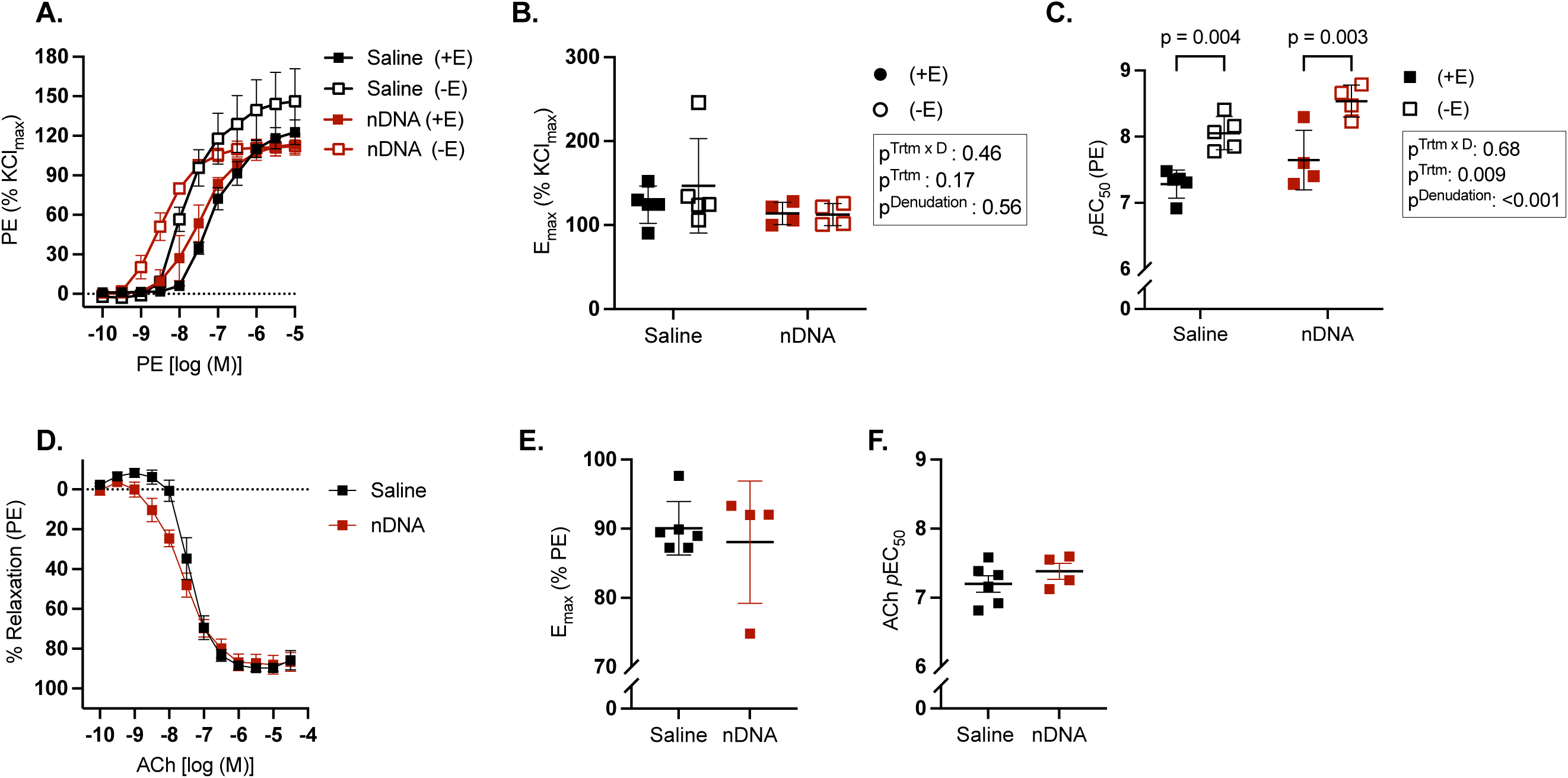
Aortic contractile and relaxation responses after in vivo saline or nuclear DNA (nDNA) challenge. Cumulative concentration-response curves (CCRCs) and corresponding maximum responses (E_max_) and pEC_50_ values to phenylephrine (PE; A-C) and acetylcholine (ACh; D-F) were assessed in thoracic aortic rings from pregnant rats 4 h after intravenous injection of saline (black symbols) or nDNA (red symbols). Open symbols show the assays in endothelium-denuded (-E) aorta rings and closed symbols in intact (+E) aorta rings. E_max_ (B, E) represents the maximum contractile or relaxation response and pEC_50_ (C, F) is the negative logarithm of the EC_50_. Data were analyzed by two-way ANOVA with Sidak post hoc test. CCRCs are presented as mean ± SEM. pEC_50_ and E_max_ values are shown as individual data points with bars representing mean ± SD; n=4-5 rats/group. For these comparisons, the saline control group was derived from the corresponding saline dataset presented in Figures 2, 3.

#### In vivo mtDNA challenge – uterine artery reactivity

To determine whether the effects of mtDNA were vascular bed specific, we also assessed dilatory and contractile responses in uterine arteries from pregnant rats treated with mtDNA or saline. ACh-induced relaxations did not differ between groups (E_max_, p=0.29, Mann-Whitney *U* test; pEC_50_, p=0.54, unpaired t-test; all n=6-7 rats/group; **Figure S3**). Similarly, SNP-induced relaxations were comparable between saline and mtDNA groups (E_max_, p=0.18, Mann-Whitney *U* test; pEC_50_, p=0.19, unpaired t-test; all n=6-7 rats/group; **Figure S3**). KCl-mediated uterine artery constrictions were also similar between groups (AUC, p=0.14, unpaired t-test; E_max_, p=0.23, unpaired t-test; all n=6-7 rats/group; **Figure S3**). These data indicate that despite a systemic inflammatory response and endothelial impairments in pregnant aortas, uterine artery function was not affected by exposure to mtDNA in pregnant rats.

#### In vivo TLR9 antagonism

To assess the contribution of TLR9 to impaired ACh-mediated relaxations in pregnant rats, we studied a separate cohort, in which pregnant rats received intravenous injections of saline, mtDNA, ODN2088, or ODN2088+mtDNA (**Figure 5A-B)**. We observed a decrease in ACh pEC_50_ in the mtDNA group compared with saline (one-way ANOVA, F(3,19) = 5.645, p=0.0061; Tukey’s post hoc test, saline vs. mtDNA: p=0.0051). Post hoc multiple comparisons tests demonstrated that ACh pEC_50_ was comparable between mtDNA and ODN2088+mtDNA groups (p=0.24), and no differences were observed between saline and ODN2088+mtDNA (p=0.15) or between saline and ODN2088 (p=0.62) (5-7 rats/group; **Figure 5A-B**). These data suggest that pharmacological antagonism of TLR9 partially reduced mtDNA-induced maternal endothelial dysfunction, as ODN2088+mtDNA-treated rats did not differ significantly from saline controls, yet the mtDNA and ODN2088+mtDNA groups also did not differ significantly from each other.

**Figure 5.**
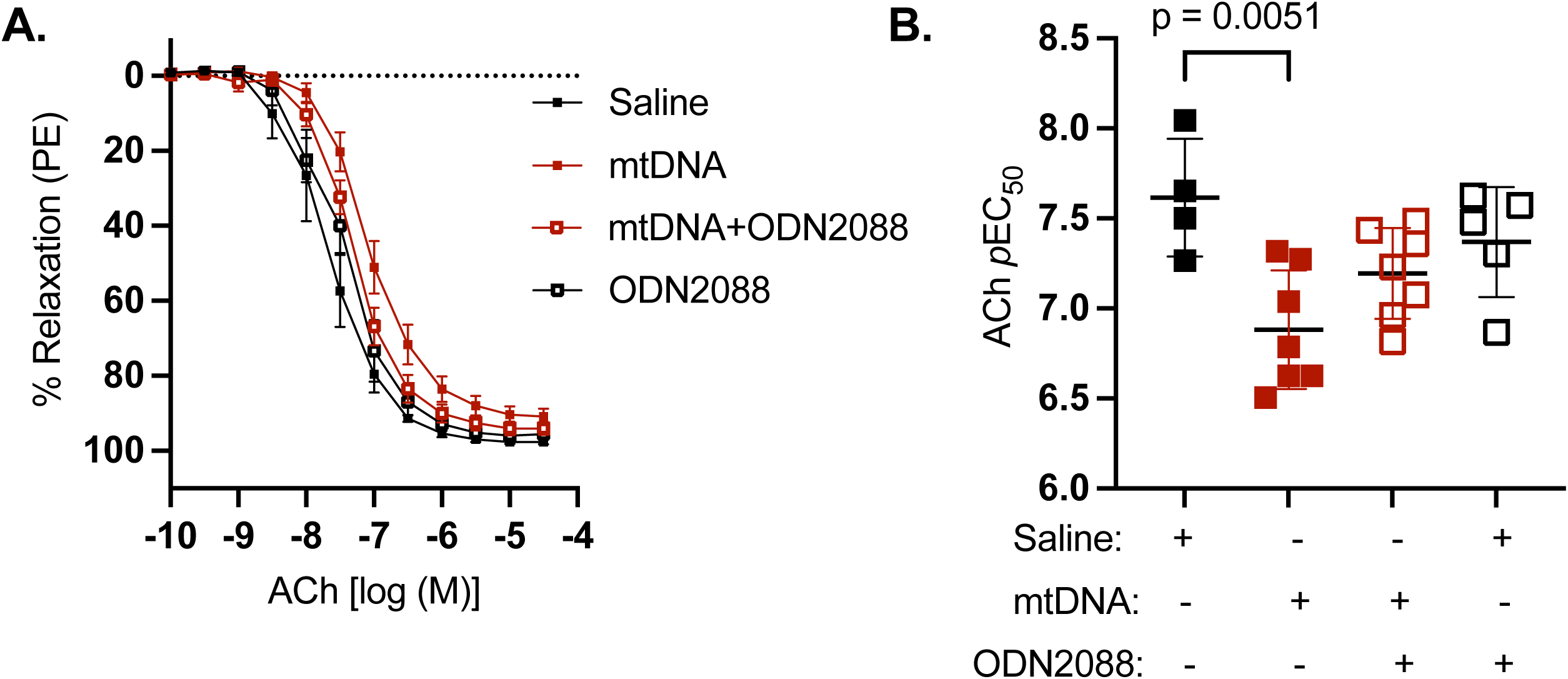
Aortic endothelium-dependent relaxation responses in pregnant rats following in vivo mitochondrial DNA (mtDNA) challenge with or without toll-like receptor 9 (TLR9) antagonism. Cumulative concentration-response curves to acetylcholine (ACh) and corresponding pEC_50_ values were assessed in thoracic aortic rings from pregnant rats 4 h after intravenous injection of saline (black symbols) or mtDNA (red symbols) in the absence (closed symbols) or presence (open symbols) of the TLR9 antagonist ODN2088. ODN2088 was administered intravenously 5 min before mtDNA or saline treatment. pEC_50_ is the negative logarithm of the EC_50_. Data were analyzed by one-way ANOVA with Tukey’s post hoc test. CCRCs are presented as mean ± SEM. pEC_50_ values are shown as individual data points with bars representing mean ± SD; n=4-7 rats/group.

Because the magnitude of the mtDNA-induced effect was smaller in this cohort than in Study 1, we examined whether inter-cohort differences in animal characteristics or baseline vascular responses contributed to this difference. We compared maternal body weights, litter size, gestational day distribution, and ACh responses across all four groups using two-way ANOVA and Fisher’s exact test **(Table 1)**. There were no differences in litter size and gestational day distribution within the GD14-15 window between the four groups with no cohort main effect or treatment × cohort interactions (p>0.05). Although maternal body weight differed between cohorts as a main effect (p=0.04), the absence of litter size difference and gestational day distribution difference suggest that this may reflect batch variability rather than a difference in pregnancy state at the time of treatment. Two-way ANOVA revealed significant main effects of both treatment and cohort on ACH E_max_ (treatment: p=0.003; cohort: p=0.002) and ACh EC50 (treatment: p=0.003; cohort: p=0.02), with no significant treatment × cohort interactions for either outcome (p>0.05). The significant treatment main effects confirm that mtDNA consistently impaired maternal endothelium-dependent relaxation across cohorts, while the cohort main effects reflect differences in baseline vascular reactivity between Study 1 and Study 3 saline-treated groups.

**Table 1.** Maternal characteristics and baseline vascular reactivity across treatment groups in Study 1 (in vivo mtDNA challenge) and Study 3 (TLR9 antagonism) cohorts.

| Parameter | Study 1 –<br>Saline (n=6) | Study 1 –<br>mtDNA<br>(n=8) | Study 3 –<br>Saline (n=4) | Study 3 –<br>mtDNA<br>(n=7) | Treatment<br>effect | Cohort<br>effect | Interaction |
| --- | --- | --- | --- | --- | --- | --- | --- |
| **Maternal**<br>**body weight**<br>**(g)** | 285.7 ± 41.8 | 287.1 ± 47.5 | 244.1 ± 19.4 | 260.4 ± 26.1 | p=0.57 | p=0.04 | p=0.63 |
| **Litter size** | 13.0 ± 3.3 | 12.8 ± 4.8 | 12.3 ± 1.0 | 11.4 ± 4.7 | p=0.87 | p=0.55 | p=0.13 |
| **ACh E<sub>max</sub>** | 90.08 ± 3.9 | 62.09 ± 20.7 | 97.8 ± 0.95 | 91.01 ± 5.1 | p=0.003 | p=0.002 | 0.052 |
| **ACh EC<sub>50</sub>** | 7.2 ± 0.29 | 5.9 ± 1.1 | 7.6 ± 0.32 | 6.88 ± 0.33 | p=0.002 | p=0.02 | p=0.37 |
| **GD14/15, n** | 3/3 | 5/3 | 1/3 | 3/4 | Fisher's<br>exact p=0.75 |  |  |

mtDNA, mitochondrial DNA; TLR9, Toll-like receptor

#### Ex vivo mtDNA challenge

To determine whether mtDNA alters vascular function through direct interactions with the vascular wall (as opposed to downstream systemic influences), we incubated ex vivo aortic rings from pregnant rats with mtDNA (0.2, 2, or 20 ng/µL) for 4 h in the presence or absence of WBCs (1 × 10^6^ cells/vessel) prior to vascular function assessment. We observed no differences in ACh-mediated relaxation across groups (all p≥0.40; one-way ANOVA with Tukey’s multiple comparisons test; n=4 rats/group; **Figure S4).**

### Molecular and biochemical correlates of vascular dysfunction

Because mtDNA selectively impaired endothelium-dependent relaxations in aortas from pregnant rats, we next tested whether this phenotype reflected altered NO signaling and NO signaling modulators, including oxidative stress and inflammatory molecules. Key assays are shown in **Figure 6 (A-D),** with additional NO/redox and inflammatory outcomes provided in **Figures S5-S6)**.

**Figure 6.**
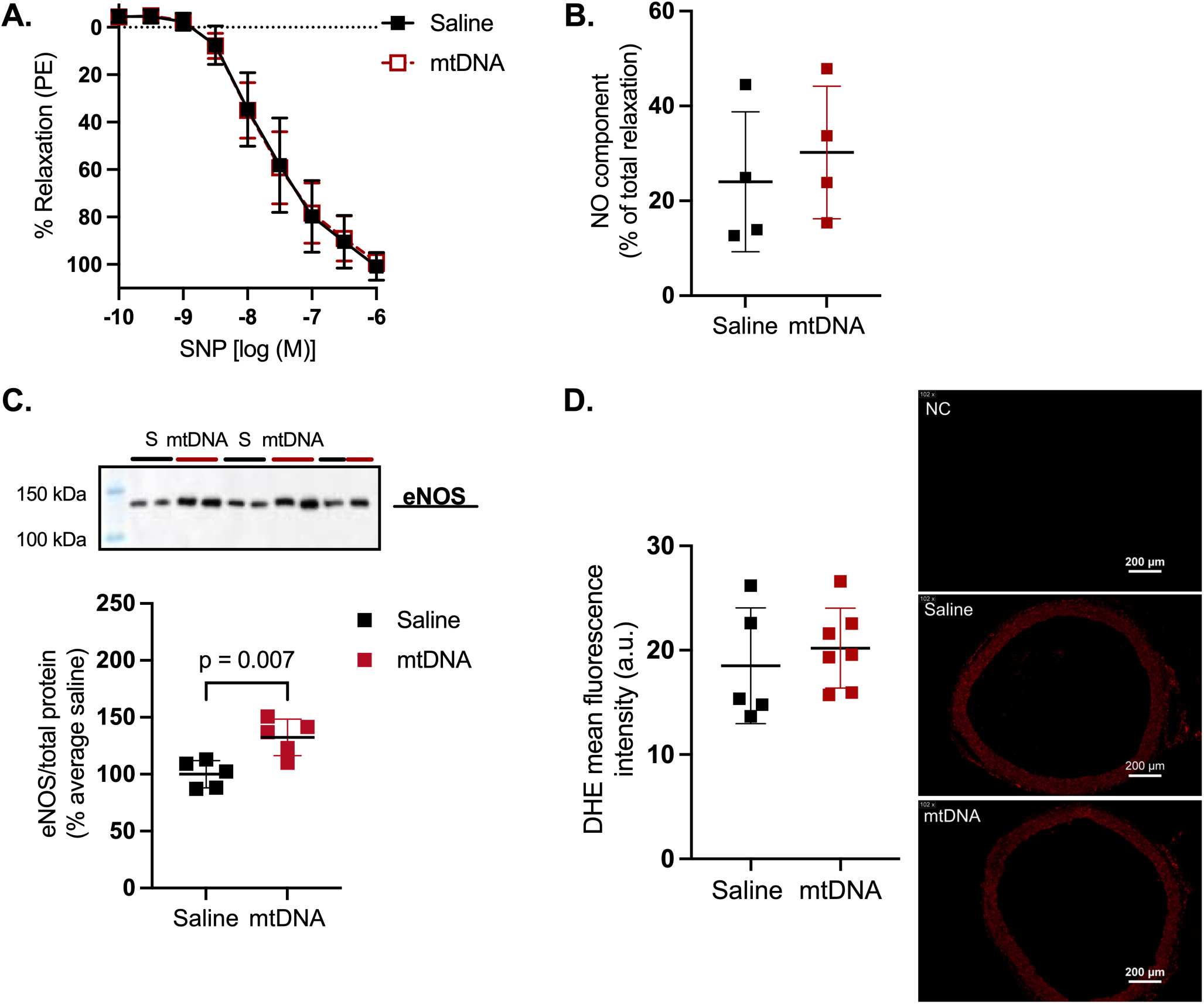
Molecular and biochemical correlates of mitochondrial DNA (mtDNA)-induced endothelial dysfunction in aortas from pregnant rats. (A) Relaxation responses to the NO donor sodium nitroprusside (SNP) were assessed in endothelium-denuded thoracic aortic rings. (B) Relative NO contribution to ACh-mediated relaxation was calculated from acetylcholine (ACh) cumulative concentration-response curves ± the NOS inhibitor L-NAME and expressed as percent NO contribution. (C) Aortic endothelial NO synthase protein content. Data were analyzed using Mann–Whitney *U* tests (B) or unpaired t-tests (C). SNP concentration-response curves are presented as mean ± SEM. The remaining data are shown as individual data points with bars representing mean ± SD. Representative image for eNOS is shown in C.

First, we sought to investigate if reduced ACh-induced relaxations reflect impaired vascular smooth muscle responsiveness to NO. Dilatory responses to NO donor SNP in endothelium-denuded aortas were comparable between saline and mtDNA groups (**Figure 6A**), suggesting preserved vascular smooth muscle sensitivity to NO. Second, we performed ACh CCRCs in the presence of a NOS inhibitor L-NAME to assess the effects of mtDNA on the relative contribution of NOS-derived NO to ACh-induced relaxations. NOS inhibition reduced ACh-induced relaxation in both saline and mtDNA-treated pregnant groups **(Figure S5A).** Accordingly, the NO contribution to ACh CCRC was comparable between groups (NO contribution to ACh CCRC (%), Median (IQR), saline: 105.7 (29.6), mtDNA: 90.70 (23.65), Mann-Whitney *U* test, p=0.20; **Figure 6B**), indicating that relative NO contribution to ACh remained unchanged following the treatment with mtDNA. We next tested predefined NO-, redox-, and inflammatory-related mechanisms that commonly modulate endothelial function. Despite reduced ACh-mediated dilation, eNOS protein abundance was increased in aortas from mtDNA-treated rats compared with saline-treated controls (p<0.007, n=5 rats/group, unpaired t-test; **Figure 6C**), suggesting potential upregulation of eNOS in response to mtDNA exposure. Plasma nitrate and nitrite concentrations (nitrate, p=0.54, Mann-Whitney *U* test, n=8-9 rats/group; nitrite, p=0.77, unpaired t-test, n=8-9 rats/group; **Figure S5B)** and DAF-FM mean fluorescence intensity in aortas were comparable between saline and mtDNA groups (DAF-FM: n=5-6 rats/group; p=0.82, unpaired t-test; **Figure S5C**), suggesting no major systemic reduction in NO bioavailability and no changes in aortic bulk NO production in response to mtDNA. Aortic DHE mean fluorescence intensity was comparable between saline and mtDNA groups (DHE: n=5-7 rats/group; p=0.55, unpaired t-test; **Figure 6D**). These results indicate an increase in eNOS abundance in aortas from pregnant rats but not downstream measurable changes in NO signaling and vascular smooth muscle sensitivity to NO in response to mtDNA exposure.

Aortic *sod-1* mRNA expression was decreased in mtDNA-treated pregnant rats relative to saline controls (p=0.0358, n=4-5 rats/group, unpaired t-test), while there were no group differences in *sod-2 and catalase mRNA,* or SOD-1, and Catalase protein abundance (all p>0.05; **Figure 7).** SOD-2 protein content increased in aortas from mtDNA-treated pregnant rats (p=0.0556, n=5 rats/group, Mann-Whitney *U* test). Aortas from mtDNA-treated rats had reduced *tnf-α* expression compared to saline-treated controls (n=4-5 rats/group; p=0.008, unpaired t-test; **Figure S6A**), whereas *mcp-1* and *il-1β* mRNA levels were unchanged (n=4-6 rats/group; p≥0.25, unpaired t-test; **Figure S6B-C)**. All immunoblots are presented in supplementary materials **(Figures S7-S10).**

**Figure 7.**
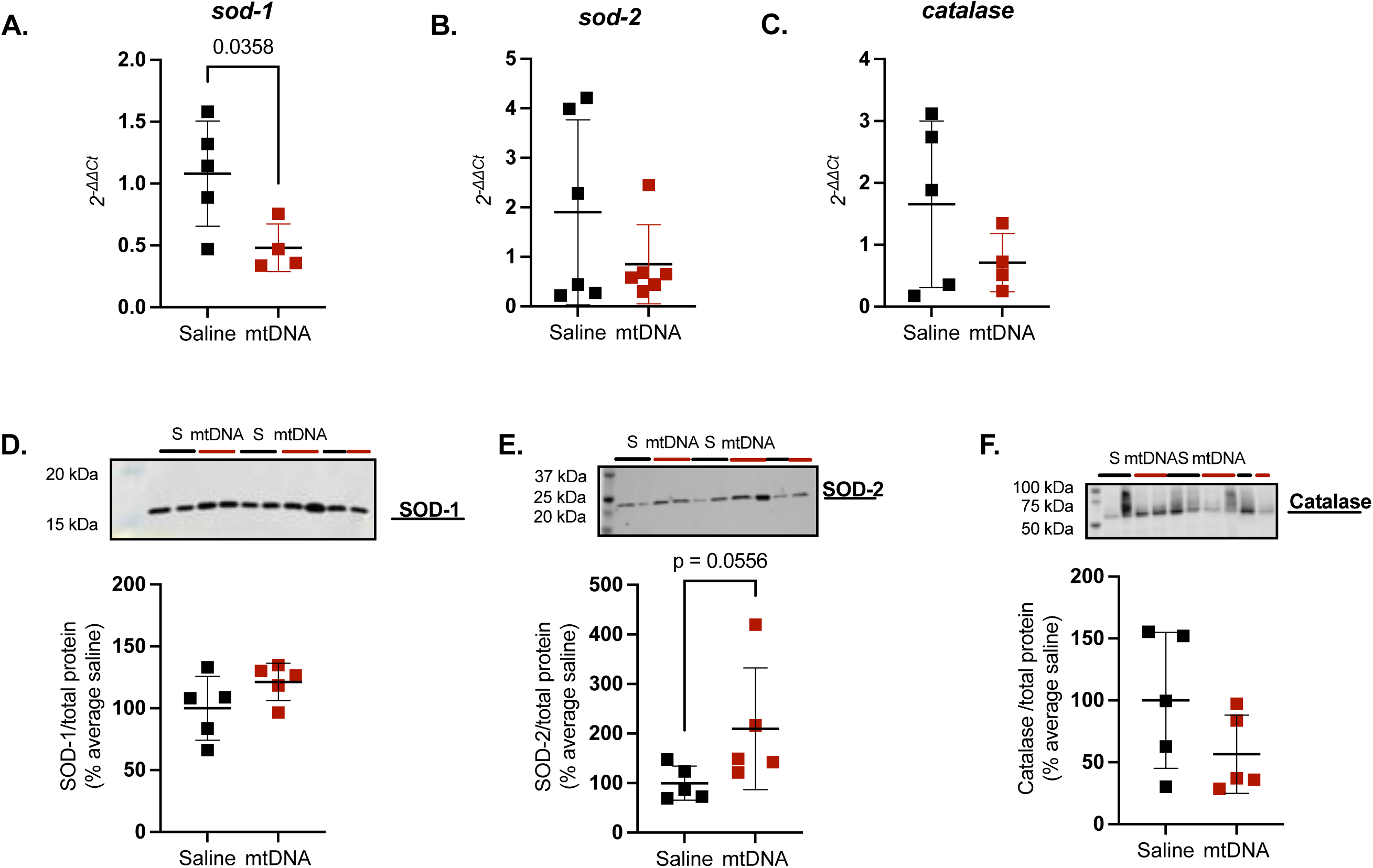
Antioxidant gene and protein expression in aortas from pregnant rats following in vivo saline or mitochondrial DNA (mtDNA) challenge. Thoracic aortas were collected from pregnant rats 4 h after intravenous injection of saline (control) or mtDNA. (A-C) Aortic mRNA expression of *sod-1* (A; n=4-5 rats/group), *sod-2* (B; n=6 rats/group), and *catalase* (C; n=4-5 rats/group). (D-F) Aortic protein content of SOD-1 (D), SOD-2 (E), and Catalase (F), with representative immunoblots shown for each protein (all n=5 rats/group). For mRNA analyses, statistical testing was performed on ΔCt values. Data were analyzed using unpaired t-tests (A, D) or Mann–Whitney *U* tests (B, C, E, F). Data are shown as individual data points with bars representing mean ± SD.

Collectively, these data demonstrate that the mtDNA-induced impairment of endothelium-dependent relaxation is not explained by impaired smooth muscle NO responsiveness, NOS-dependent vasodilation, or overt vascular oxidative/inflammatory activation at this timepoint.

## Discussion

In the present study, we evaluated the inflammatory and vasoactive potential of purified mtDNA and determined how pregnancy modifies these responses. Acute mtDNA exposure induced early innate immune transcriptional activation in the liver and lung of both non-pregnant and pregnant rats, but the magnitude and pattern of cytokine gene upregulation were pregnancy- and organ-specific. Importantly, acute mtDNA exposure induced endothelial dysfunction in the aorta of pregnant but not non-pregnant rats, while uterine artery function was preserved, indicating vascular bed-specific effects. This aortic endothelial dysfunction was partially attenuated by TLR9 antagonism and was not explained by impaired smooth muscle responsiveness to NO, NOS-dependent vasodilation, or overt vascular oxidative and inflammatory activation at this early time point. The inability to replicate the in vivo vascular effects ex vivo suggests that systemic rather than direct vascular mechanisms mediate mtDNA-induced endothelial dysfunction during pregnancy.

Extracellular mtDNA can act as a DAMP capable of activating innate immune pathways through multiple sensors, and it has been implicated in inflammatory states, including pregnancy complications (reviewed in (49)). To our knowledge, this is the first study to reveal that the magnitude of inflammatory transcriptional responses to purified mtDNA across lung and liver differ between pregnant and non-pregnant states, being relatively greater in the non-pregnant than pregnant group. These findings suggest that pregnancy shifts the immune system toward a more tolerant phenotype in response to pro-inflammatory stimuli such as mtDNA, in an organ- and pathway-specific manner, with the exception of *ifn-γ,* a pro-Th1 inflammatory marker, which was upregulated in both liver and lung of pregnant, but not non-pregnant rats.

Previous studies suggest that pregnancy modulates the expression of cytokines and associated signaling pathways that contribute to systemic tolerance and a local immunosuppressive state in various organs such as the liver (reviewed in (50)). For instance, Yang et al. showed a downregulation of key inflammatory cytokines such as IFNγ, IL-2, IL-4, IL-6 and IL-10 in the liver of pregnant ewes compared to non-pregnant controls (51). A potential factor that may be involved in the regulation of hepatic responses to pro-inflammatory stimuli is changes in the maternal hormonal profile during pregnancy. These include increased estrogen and progesterone levels (52–54) that could promote regulatory immune pathways and are associated with reduced inflammatory cytokine signaling in the maternal liver. These pregnancy-associated hormonal changes may therefore contribute to the attenuated hepatic transcriptional response to mtDNA observed in pregnant compared to non-pregnant rats in the current study.

In contrast to the broadly attenuated hepatic response, the pulmonary transcriptional response to mtDNA during pregnancy showed a more selective pattern of activation. We showed an increased expression of pro-inflammatory *tnf-α* and *ifn-γ* in the lung following mtDNA exposure during pregnancy, whereas the expression of several other cytokines was reduced or unchanged. Notably, lung *il6* expression was decreased in pregnant rats following mtDNA treatment. This finding was unexpected given the well-established role of IL-6 as a proinflammatory mediator in preeclampsia and other pregnancy complications (55). This reduction may reflect pregnancy-specific regulatory mechanisms that limit IL-6 signaling in the pulmonary compartment, potentially involving IL-6 trans-signaling modulation or the anti-inflammatory actions of pregnancy hormones or pulmonary immune cells (56); however, the precise mechanism warrants further investigation.

Pregnancy is associated with substantial structural, functional, and immunological changes in the pulmonary system (57). Vermillion *et al.* reported that pregnant female mice exhibit higher lung compliance, total lung capacity, and fixed lung volume compared with non-pregnant controls, and that these physiological changes help preserve pulmonary function during influenza infection (58). These findings suggest that pregnancy-induced changes in lung mechanics create a physiologically distinct pulmonary environment that modifies the organ’s response to immune challenges. By extension, similar pregnancy-induced adaptations in pulmonary structure and immune tone may determine the lung transcriptional response to sterile immune stimuli such as circulating mitDNA, contributing to selective cytokine pattern observed in our study. In addition, pregnancy alters susceptibility and severity of certain respiratory infections. Pneumonia may be more severe during pregnancy due in part to circulatory changes and reductions in functional residual lung capacity caused by increased abdominal pressure (3, 59), further underscoring that the pregnant lung operates under a different physiological baseline. The selective upregulation of *tnf-α* and *ifn-γ* in the lungs of pregnant rats followed mtDNA exposure may reflect a shift toward innate antiviral-type immune surveillance in the pulmonary compartment during pregnancy, consistent with the known role of these cytokines in respiratory host defense.

A key and novel finding of this study was the pregnancy-specific impairment of endothelium-dependent relaxation in aortas from pregnant but not non-pregnant rats. The contribution of the endothelium to PE-induced vasoconstriction was also impaired during pregnancy following mtDNA exposure, while overall aortic contractile capacity and smooth muscle sensitivity to NO were preserved. These findings demonstrate that acute systemic mtDNA exposure is sufficient to induce aortic endothelial dysfunction in a pregnancy-specific manner.

Because endothelial function is regulated by interactions between NO signaling, redox balance, and inflammatory pathways, we examined components of these axes based on our previous work (20). Although we did not detect overt increases in vascular ROS production or changes in NO bioavailability at this early time point, we observed transcriptional changes in antioxidant pathways, including reduced *sod-1* expression and a trend toward increased *sod-2*, alongside increased aortic eNOS protein abundance. Key inflammatory cytokines did not increase in the aorta in response to mtDNA. Taken together, these molecular findings are correlates of the observed endothelial dysfunction whose functional significance remains to be determined. The absence of overt oxidative stress or NO depletion at 4 hours suggests that the endothelial impairment may precede the development of detectable redox or inflammatory changes, consistent with an early and dynamic perturbation of endothelial homeostasis rather than established oxidative or inflammatory injury. These findings contrast with our prior study, in which repeated stimulation of TLR9 with a synthetic agonist induced vascular oxidative stress and impaired NOS-dependent mechanisms in mesenteric arteries (20), suggesting that the temporal and mechanistic profile of vascular injury differs between acute purified mtDNA exposure and chronic synthetic TLR9 stimulation.

Circulating cell-free mtDNA levels were not elevated at 4 hours post-injection, consistent with the known rapid clearance kinetics of circulating DNA (60) (61) and possible cellular uptake by immune or peripheral cells. These observations suggest that even transient exposure to circulating mtDNA may be sufficient to initiate inflammatory and vascular responses that persist beyond the period of detectable plasma increases.

The mtDNA-induced upregulation of *tlr9* in the lung and liver of both pregnant and non-pregnant states is consistent with engagement of CpG-sensing pathways, and pharmacological antagonism of TLR9 partially reduced the endothelial dysfunction, supporting a role for TLR9 in mediating the vascular effects of mtDNA during pregnancy. The incomplete reversal by TLR9 antagonism suggests that additional DNA-sensing pathways may contribute to the observed vascular phenotype, including cGAS-STING or NLRP3 inflammasome (62–64). Downstream TLR9 signaling mediators such as Myeloid Differentiation Primary Response 88 (MyD88) and regulatory factor 7 (IRF7) were not assessed in the current study and warrant further investigation.

The smaller magnitude of mtDNA-induced endothelial dysfunction in Study 3 compared with Study 1 was addressed by formal between-cohort comparison of animal characteristics and baseline vascular reactivity (Table 1). Two-way ANOVA revealed consistent treatment main effects on ACh Emax and pEC50 across both cohorts with no treatment × cohort interactions, demonstrating that mtDNA impaired endothelium-dependent relaxation similarly across cohorts. The cohort main effects for ACh Emax and pEC50 reflected differences in baseline vascular reactivity between cohorts rather than differential sensitivity to mtDNA. Litter size, gestational day distribution, and maternal weight did not interact with the effect of treatment. These findings confirm the reproducibility of the mtDNA-induced endothelial impairment and support the validity of the TLR9 antagonism data.

To determine whether mtDNA acts directly on the vascular wall, we incubated aortic rings ex vivo with mtDNA with or without WBCs for 4 hours; however, vascular function remained intact under these conditions. These results suggest that the in vivo endothelial dysfunction requires systemic mediators, such as circulating immune mediators, neurohumoral signals, pr hemodynamic stimuli, rather than direct vascular actions of mtDNA.

Finally, the absence of functional changes in uterine arteries despite systemic endothelial dysfunction in the aorta demonstrates that mtDNA-mediated vascular effects are vascular bed-specific. This pattern is consistent with our previous study showing that TLR9 stimulation did not alter uterine artery function, while impairing mesenteric resistance artery responses during pregnancy (20). The uterine vasculature undergoes substantial structural and functional remodeling during pregnancy to meet fetoplacental developmental demands (35, 65), and this remodeling may confer resilience to inflammatory stimuli, though the specific mechanisms protecting uterine artery function from mtDNA-induced impairments were not directly examined in this study. Collectively, these findings support the concept that pregnancy differentially programs vascular beds, resulting in selective susceptibility of systemic vessels while preserving uterine perfusion.

### Strengths and Limitations

A major strength of the present study is the use of a controlled in vivo model with rigorous mtDNA quality control to directly assess the effects of circulating mtDNA on maternal vascular function and inflammation during pregnancy. The inclusion of nuclear DNA as a specificity control, multi-organ inflammatory assessment across liver and lung, vascular bed comparisons between aorta and uterine arteries, and pharmacological TLR9 antagonism collectively provide a mechanistically informative dataset linking a physiologically relevant DAMP stimulus to maternal endothelial dysfunction.

Several limitations should be considered. First, our experimental design involved a single systemic injection of mtDNA at one gestational time point within mid-gestation, which does not recapitulate the chronic inflammatory environment that characterizes pregnancy complications such as preeclampsia. Second, estrous cycle was not controlled in non-pregnant rats, which may have introduced variability in vascular and inflammatory outcomes. Third, donor pregnancy status was matched to recipient pregnancy status for mtDNA preparation, which introduces the possibility that biological differences between pregnant-and non-pregnant-derived mtDNA contributed in part to the observed pregnancy-specific effects. Fourth, the ex vivo negative results implicate systemic mechanisms in the in vivo vascular phenotype, the specific circulating mediators or processes responsible were not identified. Fifth, only cytosolic ROS were assessed. Mitochondrial ROS, which may independently influence vascular function, were not directly measured. Finally, our study used exogenous purified mtDNA rather than endogenously released mtDNA, and the extent to which these findings reflect the pathophysiology of conditions involving endogenous sterile DAMP release warrants further investigation.

## Conclusions

In summary, our findings demonstrate that acute exposure to circulating mtDNA during pregnancy induces systemic maternal inflammation and impairs maternal endothelium-dependent vasodilation in a pregnancy- and vascular bed-specific manner. These vascular changes are associated with transcriptional changes in antioxidant pathways and increased eNOS protein abundance in the aorta, with partial involvement of the TLR9 pathway. Because pregnancy complications such as preeclampsia are characterized by heightened inflammatory stress, these findings provide novel evidence that non-bacterial inflammatory stimuli can adversely affect maternal vascular function during pregnancy. Improved understanding of how mitochondrial danger signals influence maternal vascular physiology may help inform strategies aimed at protecting maternal cardiovascular health during complicated pregnancies.

## Supplemental material

Supplemental Tables: Table S1, Table S2, Table S3, Table S4, Table S5, Table S6

Supplemental Figures: Figure S1, Figure S2, Figure S3, Figure S4, Figure S5, Figure S6

## Acknowledgments

We thank Loma Linda University Core Facility for providing access to cryostat, Western blot imaging, and qPCR instrumentation.

## Grants

This study was supported by NIH R01 HL146562 (SG), AHA 24POST1198395 (NH), AHA 24POST1198627 (RNOS), and the Dean’s Stipend Award from the School of Medicine, Loma Linda University (DE).

## Disclosures

No conflicts of interest, financial or otherwise, are declared by the author(s).

## Disclaimers

The content is solely the responsibility of the authors and does not necessarily represent the official views of the National Institutes of Health.

## Author Contributions

NH and SG conceived and designed research; NH, RNOS and SG analyzed data; NH, SG, RNOS, LL, TL, IG, GK, MR performed experiments; NH, RNOS, TL, EMG, AB, NRP, XQH, LZ and SG interpreted results of experiments; NH and SG prepared figures; NH and SG drafted manuscript; NH, RNOS, DE, LL, TL, EMG, AB, XQH, LZ and SG edited and revised manuscript; NH, RNOS, DE, LL, TL, IG, GK, MR, EMG, AB, NRP, XQH, LZ and SG approved final version of the manuscript.

## Notes

### Competing Interest Statement

The authors have declared no competing interest.

