## Supplementary Materials for "Acute exposure to cell-free mitochondrial DNA induces pregnancy-specific aortic endothelial dysfunction and organ-selective inflammation in rats"

**Affiliations:** [1]Lawrence D. Longo, MD Center for Perinatal Biology, Department of Basic Sciences, Loma Linda University School of Medicine, Loma Linda, California, USA; [2]Department of Gynecology and Obstetrics, Loma Linda University School of Medicine, Loma Linda, California, USA; [3]Department of Pediatrics, Division of Neonatology, Loma Linda University School of Medicine, Loma Linda, CA, USA; [4]Department of Microbiology, Immunology, and Genetics, University of North Texas Health Science Center, Fort Worth, Texas, USA

### Table S1. Chemicals and reagents.

| Chemical / Reagent Name | Catalog No. | Vendor/City, State, Country |
|---|---|---|
| Acetylcholine chloride | A6625 | Sigma Aldrich, Saint Louis, MO, USA |
| Antibiotic-Antimycotic | 15-240-062 | Gibco, Billings, MT, USA |
| $\beta$-mercapoethanol | M3148-25 ML | Sigma Aldrich |
| Bovine Serum Albumin | A6003-25G | Sigma Aldrich |
| Calcium chloride dihydrate | C3881 | Sigma Aldrich |
| Chromogenic Endotoxin Quant Kit | A39552 | Thermo Fisher Scientific, Waltham, MA, USA |
| DAF-FM diacetate | D23844 | Invitrogen, Carlsbad, CA, USA |
| Dextrose anhydrous | BP350-1 | Thermo Fisher Scientific |
| Dihydroethidium | 37291 | Sigma-Aldrich |
| DMEM (Dulbecco's Modified Eagle's Medium) | 17-205-CV | Corning, Corning, NY, USA |
| DNeasy Blood & Tissue Kit | 69504 | Qiagen, Redwood City, CA, USA |
| DreamTaq PCR Master Mixes (2X) | K1081 | Thermo Fisher Scientific |
| Extract-N-AMP-Tissue PCR Kit | XNAT2-1KT | Sigma Aldrich |
| Extraction Solution | E7526-24ML | Sigma Aldrich |
| Fetal Bovine Serum | A5670401 | Gibco |
| Fetal Bovine Serum, charcoal stripped | A3382101 | Gibco |
| iQ SYBR Green Supermix | 1708882 | Bio-Rad, Hercules, CA, USA |
| Isoflurane | 200-237 V1 502017 | Vet One Fluriso, Union City, CA, USA |
| L-glutamine | 25030081 | Gibco |
| Magnesium sulfate heptahydrate | M2773 | Sigma Aldrich |
| miRNeasy Mini Kit | 217004 | Qiagen |
| Mitochondria Isolation Kit | 89801 | Thermo Fisher Scientific |
| $Na_2H_2P_2O_7$ (Sodium pyrophosphate dibasic, 10 mM) | 71501 | Sigma-Aldrich |
| $Na_3VO_4$ (Sodium orthovanadate, 100 mM) | S6508 | Sigma-Aldrich |
| NaF (Sodium fluoride, 100 mM) | 201154 | Sigma-Aldrich |
| Neutralization Solution B | N3910-24ML | Sigma Aldrich |
| Nitrocellulose membrane, 0.45um 30 cm x 3.5 m | 162-0115 | Bio-Rad |
| Non-Fat Milk (Blotting-Grade Blocker) | 1706404 | Bio-Rad |
| Nω-Nitro-L-arginine methyl ester hydrochloride | N5751 | Sigma-Aldrich |
| Oligo-dT primers, 100 ul (0.4 ug/ul) | 79237 | Qiagen |
| Penicillin-Streptomycin | 15140122 | Gibco |
| Phenylephrine hydrochloride | P6126 | Sigma-Aldrich |
| PMSF (Phenylmethanesulfonyl fluoride, 100 mM) | 36978 | Thermo Fisher Scientific |
| Potassium chloride | P9541 | Sigma Aldrich |
| Potassium phosphate monobasic | P0662 | Sigma Aldrich |
| Protease Inhibitor Cocktail Tablets | P8340-1ML Lot#50-165-7349 | Sigma Aldrich |
| QIAamp DNA Mini Kit | 51304 | Qiagen |
| QIAzol Lysis Reagent | 57506291 | Qiagen |
| RiboGuard RNase inhibitor | E0126-40D7 | LGC Biosearch Technologies, Middleton, WI, USA |
| RPMI 1640 | 10-040-CV | Corning |
| Senscript RT Kit Reagents | 205213 | Qiagen |
| Sodium bicarbonate | S233-3 | Thermo Fisher Scientific |
| Sodium chloride | S271-3 | Thermo Fisher Scientific, Waltham, MA, USA |
| Sodium nitroferricyanide(III) dihydrate | 228710 | Sigma-Aldrich |
| T-PER™ Tissue Protein Extraction Reagent | 78510 | Thermo Fisher Scientific |
| TaqMan Universal Master Mix II, no UNG | 4440040 | Applied Biosystems, Waltham, MA, USA |
| Tissue Preparation Solution | T3073-3ML | Sigma Aldrich |
| Tissue-Tek O.C.T Compound | 4583 | Sakura, Torrance, CA, USA |
| Total Protein Stain | 92611011 | LI-COR Biosciences, Lincoln, NE, USA |
| Trypan Blue solution | T8154 | Sigma |
| U46619 | 16450 | Cayman Chemical, Ann Arbor, MI, USA |

**Table S2. Primer and probe sequences used for qPCR quantification of mitochondrial DNA (mtDNA) and nuclear DNA (β-actin).**

| **Target** | **Oligonucleotide** | **Sequence (5' à 3')** | **Final Concentration (µM)** | **Volume per Reaction (µL)** |
|---|---|---|---|---|
| mtDNA D-loop | Forward primer | GGTTCTTACTTCAGGGCCATCA | 0.625 | 2 |
| mtDNA D-loop | Reverse primer | GATTAGACCCGTTACCATCGAGAT | 0.625 | 2 |
| mtDNA D-loop | Probe | 6FAM-TTGGTTCATCGTCCATACGTTCCCCTTA-TAMRA | 2.5 | 1 |
| nDNA β-actin | Forward primer | GGGATGTTTGCTCCAACCAA | 2.5 | 2 |
| nDNA β-actin | Reverse primer | R: GCGCTTTTGACTCAAGGATTTAA | 2.5 | 2 |
| nDNA β-actin | Probe | VIC-CGGTCGCCTTCACCGTTCCAGTT-TAMRA | 2.5 | 1 |

**Table S3. Primer sequences used for quantitative real-time PCR analysis of gene expression in rat aorta.**

| Gene | Primer Sequence (5'➔3') | Amplicon Size (bp) |
|---|---|---|
| ***il-4*** | ***Forward:*** *CGTGATGTACCTCCGTGCTT*<br>***Reverse:*** *ATTCACGGTGCAGCTTCTCA* | 108 |
| ***il-6*** | ***Forward:*** *TGATGGATGCTTCCAAACTG*<br>***Reverse:*** *GAGCTTGGAAGTTGGGGTA* | 229 |
| ***il-1β*** | ***Forward:*** *CACCTTCTTTTCCTTCATCTTTG*<br>***Reverse:*** *GTCGTTGCTTGTCTCTCCTTGTA* | 241 |
| ***il-10*** | ***Forward:*** *CTGGCTCAGCACTGCTATGT*<br>***Reverse:*** *GCAGTTATTGTCACCCCGGA* | 86 |
| ***tnf-α*** | ***Forward:*** *ACTGAACTTCGGGGTGATTG*<br>***Reverse:*** *GCTTGGTGGTTTGCTACGAC* | 153 |
| ***mmp-8*** | ***Forward:*** *CCAAGGAGTGTCCAAGCCAT*<br>***Reverse:*** *TGCTAGTGGGGTAACCTGGA* | 127 |
| ***mcp-1*** | ***Forward:*** *CTGTCTCAGCCAGATGCAGTT*<br>***Reverse:*** *AGCCGACTCATTGGGATCA* | 80 |
| ***ifn-γ*** | ***Forward:*** *GAGGAACTGGCAAAAGGACG*<br>***Reverse:*** *CAGGTGCGATTCGATGACAC* | 133 |
| ***f4/80*** | ***Forward:*** *GCCATAGCCACCTTCCTGTT*<br>***Reverse:*** *ATAGCGCAAGCTGTCTGGTT* | 143 |
| ***sod-1*** | ***Forward:*** *TTGGCCGTACTATGGTGGTC*<br>***Reverse:*** *GGGCAATCCCAATCACACCA* | 120 |
| ***sod-2*** | ***Forward:*** *CGGGGGCCATATCAATCACA*<br>***Reverse:*** *GCCTCCAGCAACTCTCCTTT* | 84 |
| ***catalase*** | ***Forward:*** *CTGACTGACGCGATTGCCTA*<br>***Reverse:*** *ATGGTGTAGGATTGCGGAGC* | 97 |
| ***18s*** | ***Forward:*** *GCCGCTAGAGGTGAAATTCTTG*<br>***Reverse:*** *CATTCTTGGCAAATGCTTTCG* | 66 |
| ***gapdh*** | ***Forward:*** *AGACAGCCGCATCTTCTTGT*<br>***Reverse:*** *TACGGCCAAATCCGTTCACA* | 228 |

Primer sequences were designed using Primer-BLAST and validated for amplification efficiency prior to experimental analysis.

**Table S4. Primary and secondary antibodies used for Western blotting.**

| Target | Cat# | Vendor (City, State, Country) | MW (kDa) | Primary dilution | Secondary antibody (dilution; vendor; cat #) | RRIDs |
|---|---|---|---|---|---|---|
| Catalase | 14097 | Cell Signaling (Danvers, MA, USA) | 60 | 1:1000 | IRDye® 800CW Goat anti-Rabbit IgG (1:2000; LI-COR Biosciences; #926-32211) | RRID: AB_2798391 |
| SOD1 | 37385 | Cell Signaling (Danvers, MA, USA) | 16-18 | 1:1000 | Anti -Rabbit IgG, HRP-linked Antibody (1:2000; Cell Signaling Technology; #7074) | RRID: AB_3073954 |
| SOD2 | 13194 | Cell Signaling (Danvers, MA, USA) | 22 | 1:1000 | IRDye® 800CW Goat anti-Rabbit IgG (1:2000; LI-COR Biosciences; #926-32211) | RRID: AB_2750869 |
| eNOS (Clone 3) | 610297 | BD Transduction Laboratories (San Diego, CA, USA) | 140 | 1:500 | Anti-mouse IgG, HRP-linked Antibody (1:2000; Cell Signaling Technology; #7076) | RRID: AB_397691 |

**Table S5. Western blotting conditions for each target protein.**

| Protein | Cat# | Sample denaturation | % SDS gel | Electrophoresis | Transfer | Type of membrane | Blocking conditions |
|---|---|---|---|---|---|---|---|
| Catalase | 14097 | non-heated | 4-15% gradient | 2-2.5 h at 110 V | wet transfer; 120 V, 75 min | Nitrocellulose (0.45 $\mu$m) | 3% BSA (1 h) |
| SOD1 | 37385 | 95 °C (5 min) | 12% | 1 h at 80 mV and 1.5 - 2 h at 100 mV | dry transfer; 25 V, 30 min | Nitrocellulose (0.45 $\mu$m) | 3% BSA (1 h) |
| SOD2 | 13194 | non-heated | 4-15% gradient | 2-2.5 h at 110 V | wet transfer; 100 V, 90 min | PVDF | 3% BSA (1.5 h) |
| eNOS (Clone 3) | 610297 | 95 °C (5 min) | 6% | 1 h at 80 V and 1.5 - 2 h at 100 V | wet transfer; 120 V, 75 min | Nitrocellulose (0.45 $\mu$m) | 5% milk (1 h) |

Gel percentages were selected based on expected molecular weight of each target protein.

### Table S6. pEC50s, AUCs, and Emax from cumulative concentration-response curves.

| | | Non-pregnant (mean±SD) | | Pregnant (mean±SD) | | Statistical analysis | | |
|---|---|---|---|---|---|---|---|---|
| | | Saline | mtDNA | Saline | mtDNA | Int. | Preg | mtDNA |
| **ACh** | **$pEC_{50}$** | 7.55±0.09 (n=5) | 7.10±0.34 (n=6) | 7.20±0.29 (n=6) | **5.96±1.07* (n=8)** | p=0.15 | **p=0.01** | **p=0.005** |
| | **$E_{max}$** | 98.08±0.71 (n=5) | 93.44±5.30 (n=6) | 90.08±3.87 (n=6) | **62.09±20.70* (n=8)** | p=0.03 | **p=0.0008** | **p=0.004** |
| **SNP** | **$pEC_{50}$** | 8.89±0.30 (n=5) | 8.60±0.36 (n=5) | 7.64±0.64 (n=4) | 7.63±.57 (n=5) | p=0.54 | **p<0.001** | p=0.51 |
| | **$E_{max}$** | 103.7±3.30 (n=5) | 100.24±3.44 (n=5) | 100.87±11.35 (n=4) | 98.86±8.61 (n=5) | p=0.81 | p=0.46 | p=0.42 |
| **KCl** | **AUC** | 1370.57±242.67 (n=7) | 1316.50±285.70 (n=6) | 1251.46±226.79 (n=5) | 1283.97±338.94 (n=7) | p=0.71 | p=0.51 | p=0.93 |
| | **$E_{max}$** | 28.53±5.16 (n=7) | 28.63±4.90 (n=6) | 24.92±4.27 (n=5) | 25.06±6.26 (n=7) | p=0.99 | p=0.11 | p=0.96 |

| | | Non-pregnant (mean±SD) | | | | Pregnant (mean±SD) | | | | Int. | mtDNA | Denud. |
|---|---|---|---|---|---|---|---|---|---|---|---|---|
| | | Saline (+E) | Saline (-E) | mtDNA (+E) | mtDNA (-E) | Saline (+E) | Saline (-E) | mtDNA (+E) | mtDNA (-E) | | | |
| | **$pEC_{50}$** | 7.14±0.23 (n=5) | **7.99±0.06* (n=4)** | 7.21±0.24 (n=6) | **8.28±0.30* (n=5)** | | | | | p=0.29 | p=0.11 | **p<0.001** |
| | | | | | | 7.28±0.21 (n=5) | **8.05±0.25* (n=5)** | 7.47±0.56 (n=6) | 7.79±0.60 (n=5) | p=0.27 | p=0.83 | **p=0.01** |
| **PE** | **$E_{max}$** | 100.58±6.24 (n=5) | 119.33±7.29 (n=5) | 103.26±7.06 (n=6) | 129.50±16.21 (n=5) | | | | | p=0.40 | 0.15 | **p<0.0001** |
| | | | | | | 124.36±22.1 6 (n=5) | 146.95±56.2 2 (n=5) | 128.61±47.5 8 (n=6) | 120.24±9.5 5 (n=5) | p=0.38 | p=0.52 | p=0.68 |
| | **AUC** | 356.75±43.4 6 (n=6) | 447.83±137.2 0 (n=6) | 355.17±25. 71 (n=6) | 339.93±28.15 (n=6) | | | | | p=0.10 | p=0.09 | p=0.23 |
| | | | | | | 582.58±134. 22 (n=8) | 646.74±116. 48 (n=8) | 618.47±143. 81 (n=9) | 693.63±39. 29 (n=9) | p=0.89 | p=0.30 | p=0.09 |
| **U46619** | **$E_{max}$** | 130.97±17.6 5 (n=6) | 132.97±28.4 0 (n=6) | 138.09±7.68 (n=6) | **114.96±9.29# (n=6)** | | | | | p=0.99 | p=0.46 | p=0.16 |
| | | | | | | 130.64±9.63 (n=8) | **115.83±7.17# (n=7)** | 124.13±10.5 0 (n=9) | 122.14±19. 98 (n=9) | p=0.20 | p=0.90 | **p=0.06** |

* p<0.05

\# p=0.07

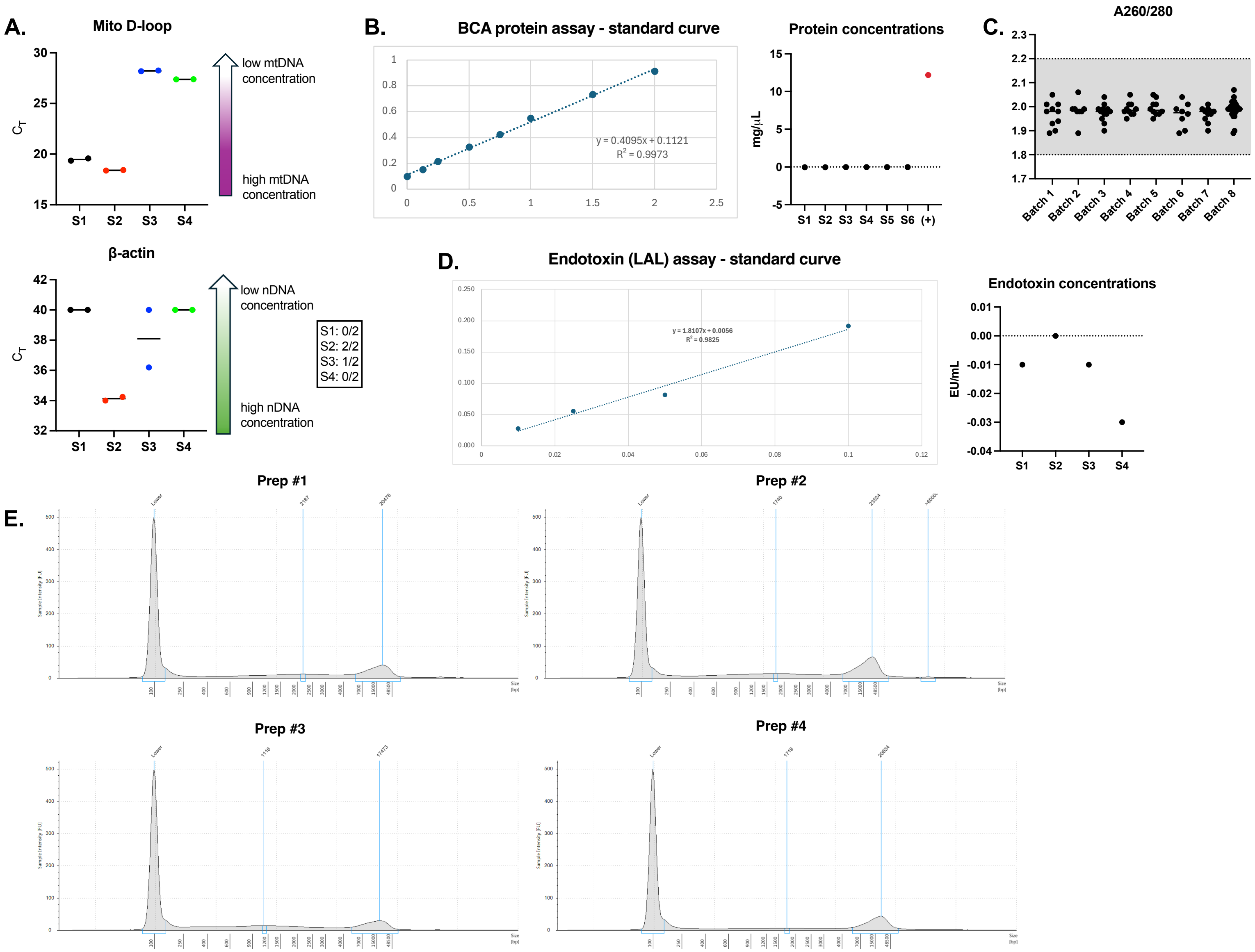

A.
Mito D-loop
CT
low mtDNA concentration
high mtDNA concentration
S1 S2 S3 S4
β-actin
low nDNA concentration
high nDNA concentration
S1: 0/2
S2: 2/2
S3: 1/2
S4: 0/2
B.
BCA protein assay - standard curve
y = 0.4095x + 0.1121
R2 = 0.9973
Protein concentrations
mg/μL
S1 S2 S3 S4 S5 S6 (+)
C.
A260/280
Batch 1 Batch 2 Batch 3 Batch 4 Batch 5 Batch 6 Batch 7 Batch 8
D.
Endotoxin (LAL) assay - standard curve
y = 1.8107x + 0.0056
R2 = 0.9825
Endotoxin concentrations
EU/mL
S1 S2 S3 S4
E.
Prep #1
Prep #2
Prep #3
Prep #4

**Figure S1. Quality control of mtDNA preparations used for in vivo studies.**

**(A)** Optimization of the mtDNA extraction protocol was assessed by TaqMan qPCR cycle threshold (Ct) values for mitochondrial D-loop (top) and nuclear β-actin (bottom) measured in four mtDNA preparations generated using protocol variants (technical replicates shown). Sample 1 exhibited the highest mtDNA yield (lowest D-loop Ct) with no detectable nuclear DNA amplification (β-actin) and was therefore selected as the final extraction protocol for subsequent studies. The inset table indicates β-actin detection rate (number of replicates with amplification/total replicates) for each sample. All reactions were run for 40 cycles. Reactions without amplification were reported as "undetermined" by instrument software and were plotted at Ct = 40 for visualization only**. (B)** Protein carryover assessment using the Pierce™ BCA Protein Assay. Left, BSA standard curve. Right, protein concentrations measured in six randomly selected mtDNA preparations. Liver lysate served as a positive control (red symbol). mtDNA preparations had protein concentrations below the assay detection threshold. **(C)** NanoDrop A260/280 ratios for mtDNA preparations across 10 isolation batches. Each point represents one preparation; shaded band indicates the acceptable range (1.8–2.1). Only preparations that were at this range were used for in vivo treatments. **(D)** Endotoxin assessment using a chromogenic Limulus Amebocyte Lysate (LAL) assay. Left, endotoxin standard curve. Right, endotoxin concentrations (EU/mL) in 4 randomly selected mtDNA preparations (and assay blank). All tested mtDNA preparations were below the assay detection threshold. **(E)** TapeStation electropherogram from four different samples showed a predominant high-molecular weight peak at ~20 kb, consistent with largely intact full-length mtDNA (expected ~16.3 kb). The arrow indicates the predominant mtDNA fragment peak. Peak broadening across samples reflects variable degrees of DNA fragmentation.

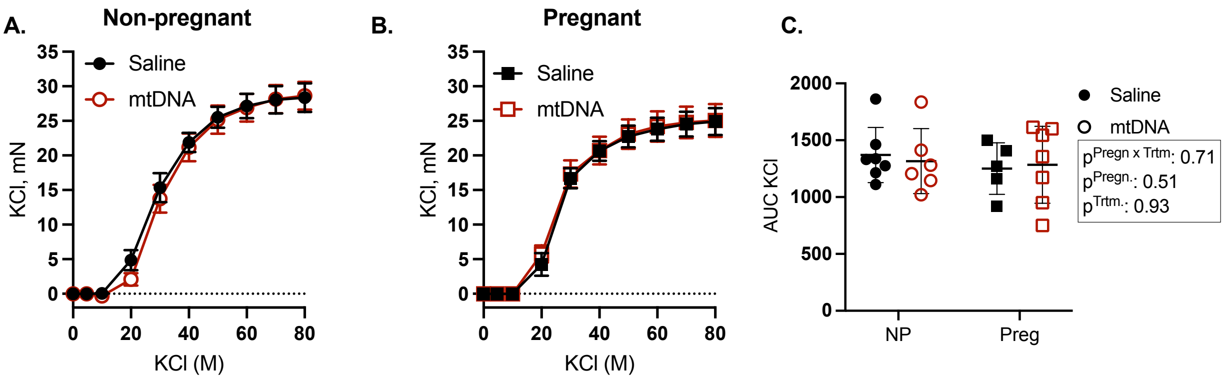


**Figure S2. High potassium-mediated constriction in rat aorta four hours after intravenous injection of purified mitochondrial DNA (mtDNA).** Cumulative concentration-response curves to potassium chloride (KCl) in aortic rings from non-pregnant (A; circle symbols) and pregnant (B; square symbols) rats. Exposure to mtDNA did not alter KCl-mediated vasoconstriction. Contractile responses are summarized as area under the curve (AUC; C; circles: non-pregnant; squares: pregnant). Concentration-response curves are presented as mean ± SEM, whereas AUC values are presented as mean ± SD with individual data points shown. Data were analyzed by two-way ANOVA with Sidak post hoc test; n = 5-6 rats/group.

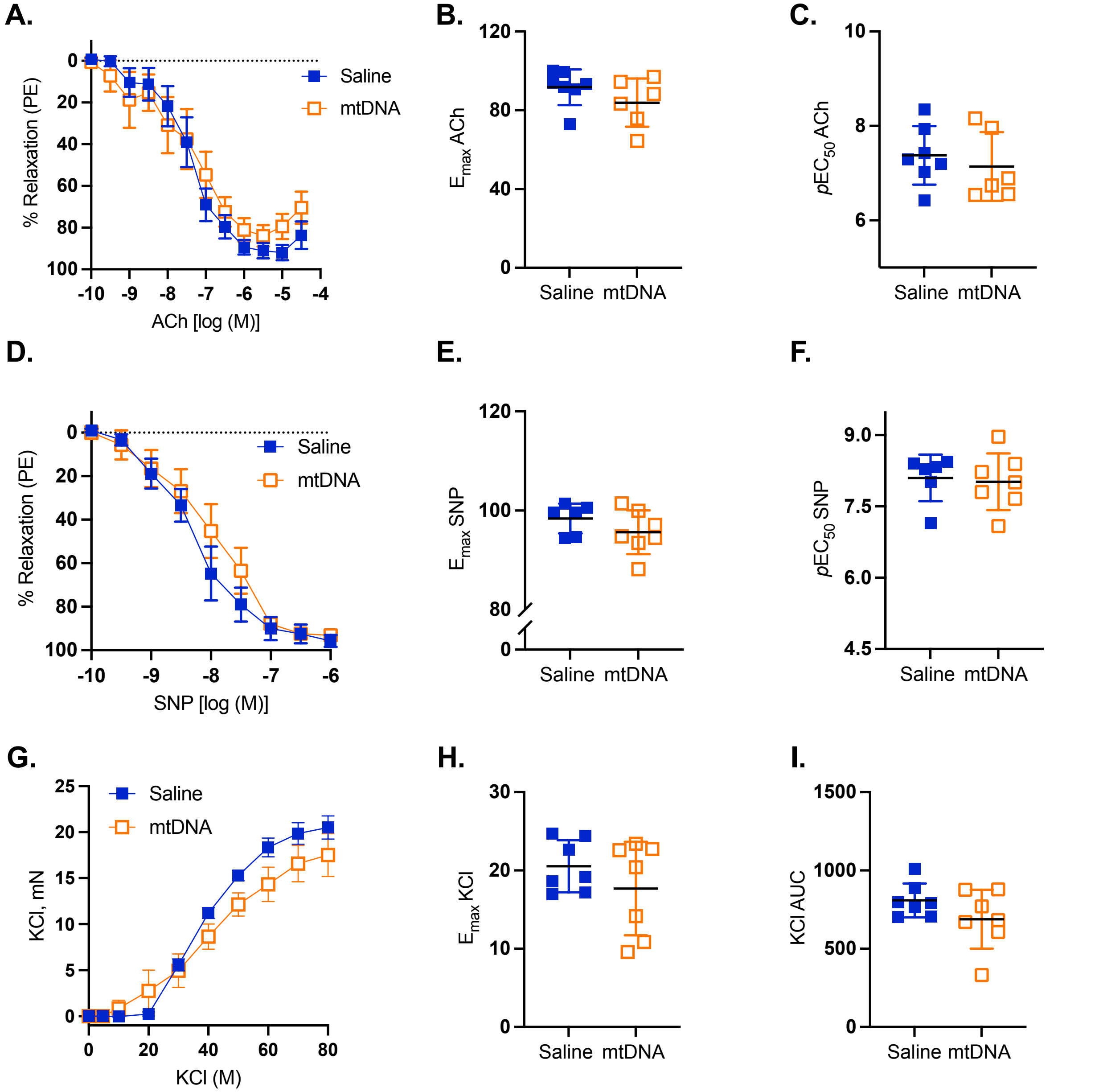


**Figure S3. Vasodilatory and contractile responses of uterine arteries from pregnant rats four hours after intravenous injection with purified mitochondrial DNA (mtDNA).** Cumulative concentration-response curves to acetylcholine (ACh; A), sodium nitroprusside (SNP, B) and potassium chloride (KCl, G) in main uterine arteries from pregnant rats. Data from saline-treated group are presented in blue, while data from mtDNA-treated group are presented in orange. Exposure to mtDNA did not affect vasodilatory responses to ACh and SNP, or contractile capacity to KCl. Vascular responses are summarized as maximal dilation ($E_{max}$; B, E, H), negative logarithm of the $EC_{50}$ ($pEC_{50}$; C, F), and area under the curve (I). Concentration-response curves are presented as mean ± SE, whereas $pEC_{50}$, $E_{max}$ and AUC values are presented as mean ± SD with individual data points shown. Data were analyzed with Mann-Whitney *U* test or unpaired t-test, as appropriate; n=5-7 rats/group.

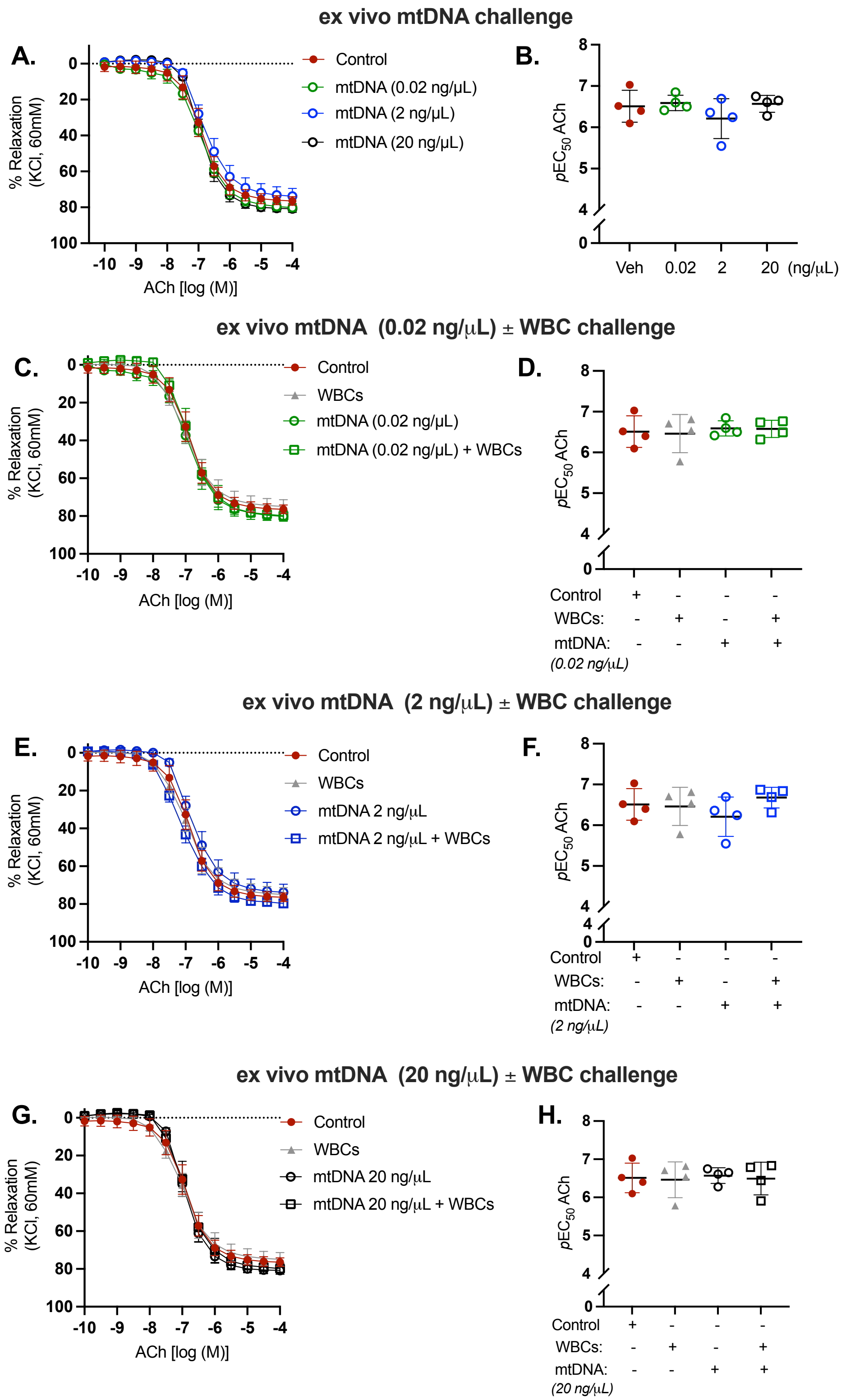


**Figure S4. Endothelium-dependent vasodilation in rat aorta four hours after in vitro incubation with purified mitochondrial DNA (mtDNA) and autologous WBCs.**

Cumulative concentration-response curves to acetylcholine (ACh) in aortic rings from pregnant rats following *in vitro* incubation with increased concentrations of mtDNA (0.02, 2, and 20 ng/$\mu$L) in the absence (A, B) or presence (C-H) of white blood cells (WBCs). Incubation with purified mtDNA, either alone or in combination with WBCs, did not alter ACh-mediated vasodilation in aorta from pregnant rats after 4 hours of *in vitro* incubation. Vasodilatory responses are summarized as the negative logarithm of the $EC_{50}$ ($pEC_{50}$; B, D, F, H). Concentration-response curves are presented as mean ± SE, whereas $pEC_{50}$ values are presented as mean ± SD with individual data points shown. Data were analyzed with one-way ANOVA with Tukey's multiple comparisons test; n=4 rats/group.

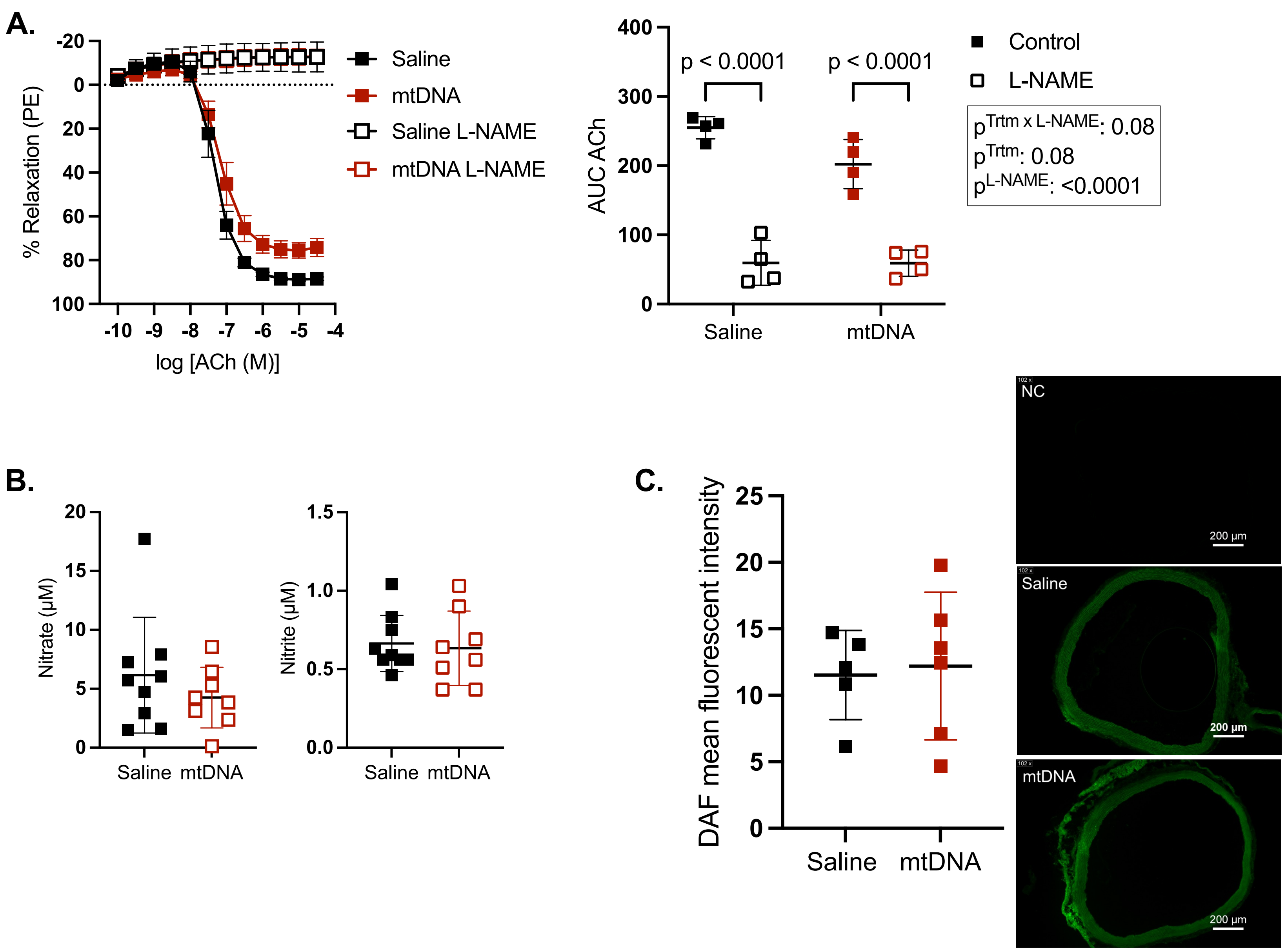
A.
% Relaxation (PE)
log [ACh (M)]
Saline
mtDNA
Saline L-NAME
mtDNA L-NAME
AUC ACh
p < 0.0001
p < 0.0001
Control
L-NAME
p^Trtm x L-NAME: 0.08
p^Trtm: 0.08
p^L-NAME: <0.0001
Saline
mtDNA
B.
Nitrate (µM)
Nitrite (µM)
Saline mtDNA
Saline mtDNA
C.
DAF mean fluorescent intensity
Saline
mtDNA
NC
Saline
mtDNA
200 µm

**Figure S5**. **Nitric oxide-related measures in pregnant rats after in vivo saline or mitochondrial DNA (mtDNA) challenge.** Thoracic aortas and plasma were collected from pregnant rats 4 h after intravenous injection of saline (black symbols) or mtDNA (red symbols). (A) Acetylcholine (ACh) cumulative concentration–response curves (CCRCs) were assessed in thoracic aortic rings in the absence (closed symbols) or presence (open symbols) of the nitric oxide synthase (NOS) inhibitor L-NAME, with corresponding area under the curve (AUC) values shown (n=4 rats/group). (B) Plasma nitrite and nitrate concentrations (n=8-9 rats/group). (C) Aortic nitric oxide-reactive fluorescence assessed by DAF-FM mean fluorescence intensity (n=5-6 rats/group). Representative fluorescence images are shown; scale bar = 200 *μ*m. Data were analyzed using two-way ANOVA with Sidak post hoc test (A), unpaired t-test (nitrite) or Mann–Whitney *U* test (nitrate) (B), and unpaired t-test (C). CCRCs are presented as mean ± SEM. AUC and fluorescence data are shown as individual data points with bars representing mean ± SD

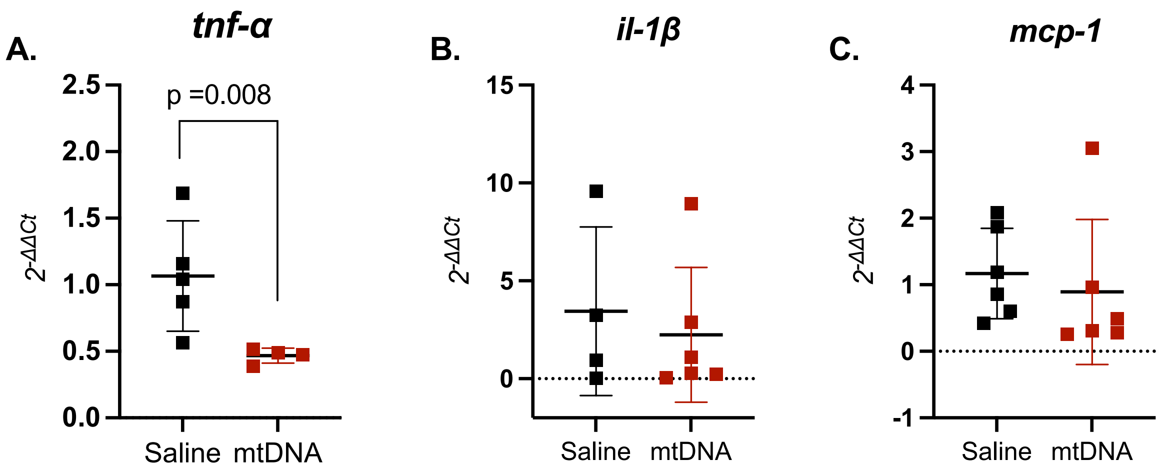


**Figure S6. Aortic inflammatory marker mRNA expression in pregnant rats following in vivo saline or mitochondrial DNA (mtDNA) challenge.** Thoracic aortas were collected from pregnant rats 4 h after intravenous injection of saline (control) or mtDNA. Relative mRNA expression levels are presented as fold change ($2^{-DDCt}$) for (A) tumor necrosis factor-α (*tnf-α*; n=4-5 rats/group), (B) interleukin 1β (*il-1β*; n=4-6 rats/group), and (C) monocyte chemoattractant protein-1 (*mcp-1;* n=6 rats/group). For mRNA analyses, statistical testing was performed on ΔCt values. Data were analyzed using unpaired t-tests (A) or Mann–Whitney *U* tests (B, C) and are presented as individual data points with bars representing mean ± SD.

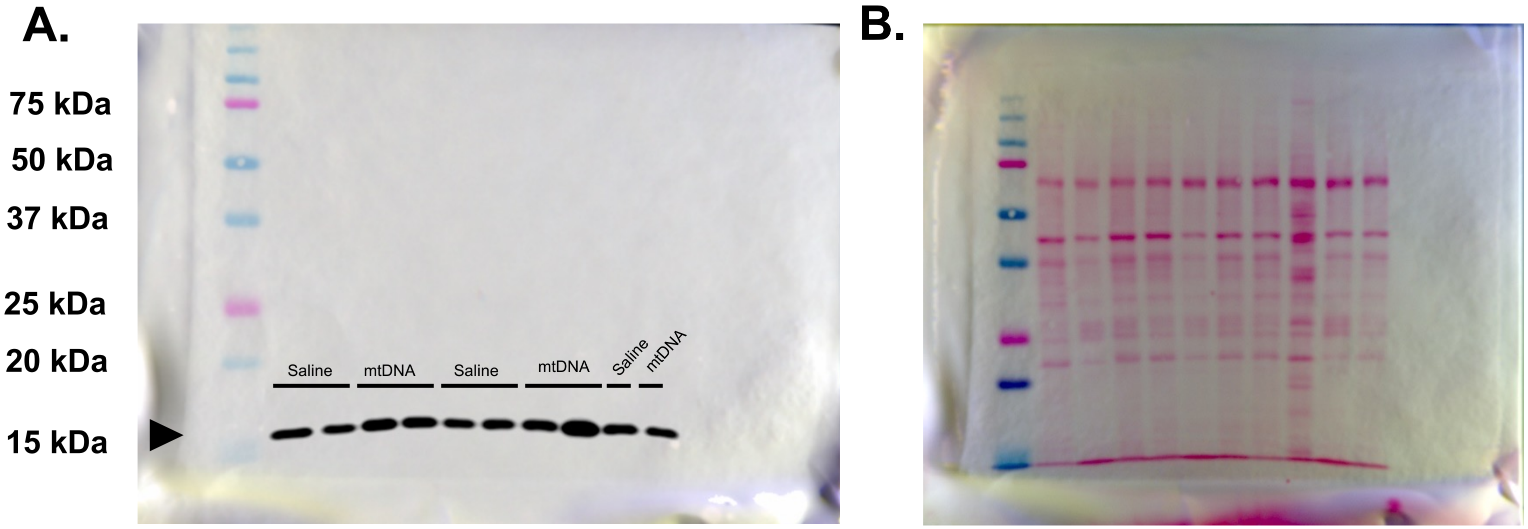


**Figure S7.** Immunoblot of superoxide dismutase 1 (SOD1, 16-18 kDa; A) and corresponding Ponceau S total protein staining (B) in aortas from pregnant rats injected with saline or mitochondrial DNA (mtDNA) on gestational days 14-15. n = 5 per group.

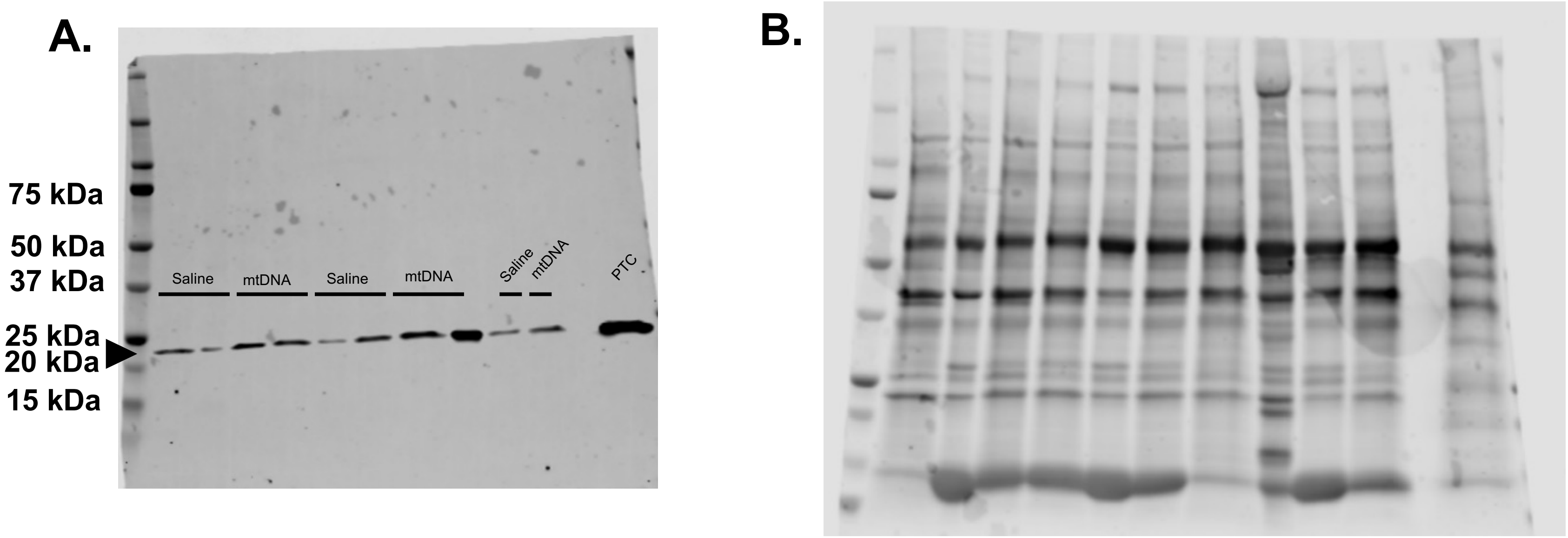


**Figure S8.** Immunoblot of superoxide dismutase 2 (SOD2, 22 kDa; A) and corresponding REVERT total protein staining (B) in aortas from pregnant rats injected with saline or mitochondrial DNA (mtDNA) on gestational days 14-15. Rat left ventricular tissue was used as a positive control (PTC). n=5 per group.

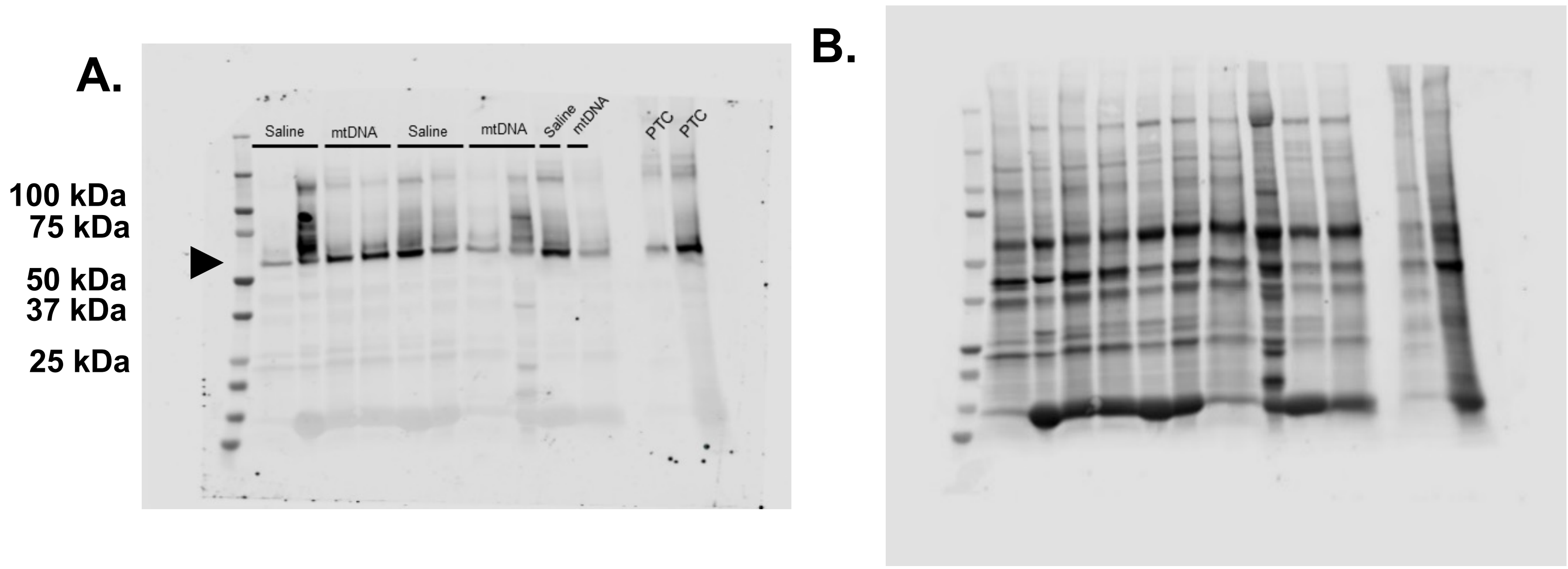


**Figure S9.** Immunoblot of Catalase (60 kDa; A) and corresponding REVERT total protein staining (B) in aortas from pregnant rats injected with saline or mitochondrial DNA (mtDNA) on gestational days 14-15. Rat left ventricular tissue was used as a positive control (PTC). n=5 per group.

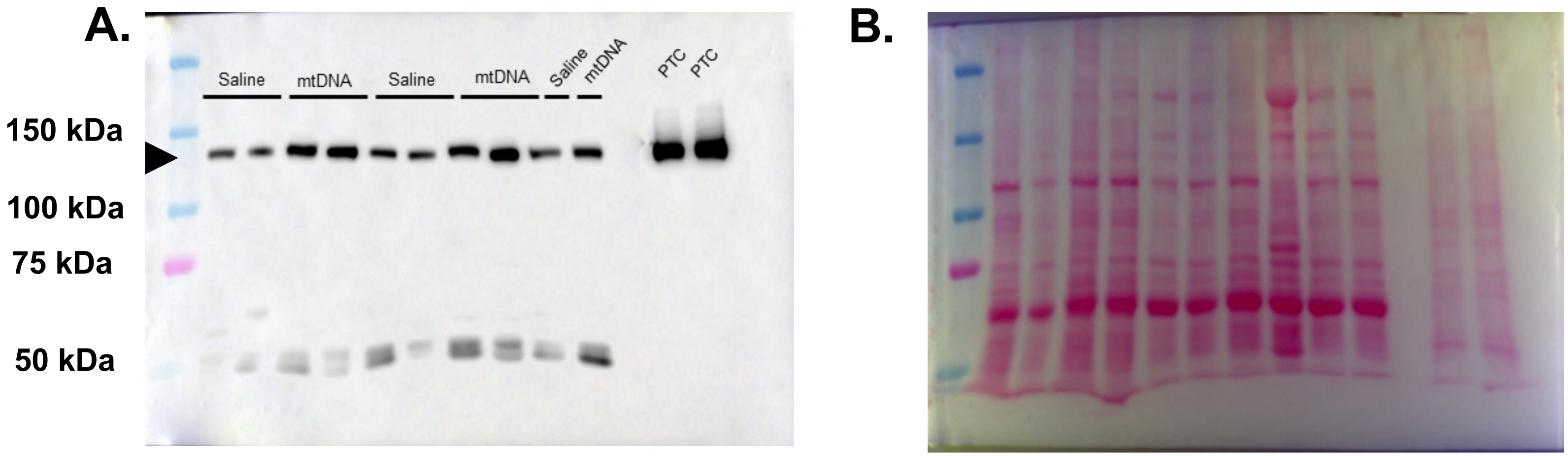


**Figure S9.** Immunoblot of endothelial nitric oxide synthase (eNOS, 140 kDa; A) and corresponding Ponceau S total protein staining (B) in aortas from pregnant rats injected with saline or mitochondrial DNA (mtDNA) on gestational days 14-15. Human umbilical vein endothelial cells (HUVECs) and HUVECs incubated with vascular endothelial growth factor (VEGF) were used as positive control (PTC). n=5 per group.
